# The mutational landscape of *Staphylococcus aureus* during colonisation

**DOI:** 10.1101/2023.12.08.570284

**Authors:** Francesc Coll, Beth Blane, Katherine Bellis, Marta Matuszewska, Dorota Jamrozy, Michelle Toleman, Joan A Geoghegan, Julian Parkhill, Ruth C Massey, Sharon J Peacock, Ewan M Harrison

## Abstract

*Staphylococcus aureus* is an important human pathogen but is primarily a commensal of the human nose and skin. Survival during colonisation is likely one of the major drivers of *S. aureus* evolution. Here we use a genome-wide mutation enrichment approach to analyse a genomic dataset of 3,060 *S. aureus* isolates from 791 individuals to show that despite limited within-host genetic diversity, an excess of protein-altering mutations can be found in genes encoding key metabolic pathways, in regulators of quorum-sensing and in known antibiotic targets. Nitrogen metabolism and riboflavin synthesis are the metabolic processes with strongest evidence of adaptation. Further evidence of adaptation to nitrogen availability was revealed by enrichment of mutations in the assimilatory nitrite reductase and urease, including mutations that enhance growth with urea as the sole nitrogen source. Inclusion of an additional 4,090 genomes from 802 individuals revealed eight additional genes including *sasA*/*sraP*, *pstA*, and *rsbU* with signals adaptive variation that warrant further characterisation. Our study provides the most comprehensive picture to date of the heterogeneity of adaptive changes that occur in the genomes of *S. aureus* during colonisation, revealing the likely importance of nitrogen metabolism, loss of quorum sensing and antibiotic resistance for successful human colonisation.

---

*Staphylococcus aureus* is an important pathogen but also commensal bacteria and part of the human microbiota. The anterior nares (lower nostrils) are regarded as the primary reservoir of *S. aureus* in humans, although the bacterium can colonise other body sites such as the skin, pharynx, axillae and perineum.^1^ Despite being a commensal, when the epithelial barrier breaks or the immune system becomes compromised, *S. aureus* can cause a variety of infections, ranging from superficial skin and soft-tissue infections to life-threatening invasive infections such as bacteraemia. Colonisation is an important risk factor for *S. aureus* infection,^2,3^ and it is frequently the strain already colonising an individual that causes an infection.^4,5^

While a few studies have sought to characterise the adaptive changes that *S. aureus* undergoes during colonisation^6,7^, our understanding remains incomplete. *S. aureus* persistently colonises ∼25% of adults, while others are either never, or only intermittently colonised.^8^ The genome of *S. aureus* encodes a range of adhesion, immune evasion and antimicrobial resistance factors that, when expressed, allow the bacterium to rapidly adapt to the nasal environment.^9–13^ In addition to changes in gene expression, mutations in the genome of *S. aureus* will also be selected during colonisation if beneficial for survival. This is supported by data from an experimental challenge model in which persistent carriers preferentially select their own strain, suggesting that *S. aureus* is adapted to the conditions on the colonised individual.^14^ This likely represents adaption to: (a) competition with other microbes in the nasal microbiota^8^; (b) nutrient availability in nasal secretions^13^; (c) adaption to the host immune response and other physiological variation; (d) spatial variation with this nasal environment (epithelium vs. hair follicles)^15^; (e) environmental exposures; and (f) the presence of therapeutic antibiotics and disinfectants (likely more acute in the clinical setting).^16^

*S. aureus* readily transmits between individuals and strain replacement may take place in persistently colonised individuals, meaning that *S. aureus* strains face common selective pressures when adapting to a new host. Mutations conferring an advantage are therefore expected to be enriched within the same genes, or groups of functionally related genes, across multiple *S. aureus* strains. To test this hypothesis, we analysed the genomes of clonal *S. aureus* isolates sampled from the same individuals, to identify evidence of adaptation in recently diverged populations of bacteria. A similar approach has recently been applied to investigate genetic changes that could promote, or be promoted by, invasive infection,^4,7,17^ or associated with persistent or relapsing *S. aureus* bacteraemia.^18–20^

To differentiate potentially adaptive genetic changes from neutral background mutation, we applied a genome-wide mutation enrichment approach to identify loci in the *S. aureus* genome under parallel and convergent evolution that could represent potential signals of adaptation during colonisation. Our results show that despite limited genetic diversity among colonising isolates of the same individual, multiple genes and pathways show a clear mutational signal of adaptation.

## Results

### Defining within-host genetic diversity in colonising isolates

To investigate putative adaptive genetic changes in *S. aureus* during colonisation we compiled a genomic dataset from 3,497 *S. aureus* colonisation isolates from ten independent studies^4,21–29^, which included a median of 2 isolates (IQR 2 to 4) from 872 individuals (Supplementary Figure 1, Supplementary Table 1). The final dataset, after excluding unrelated (non-clonal) isolates from the same host and poor-quality genomes, consisted of 3,060 isolate genomes from 791 individuals, and included 1,823 nasal isolates (59.6%), 926 isolates from multi-site screens (30.3%) and 311 isolates from other colonising sites (10.1%).

The genetic diversity between isolates colonising the same individual was low (Supplementary Figure 2), measured either as the number of single nucleotide polymorphism (SNPs) in the core genome (median 1 SNPs, IQR 0 to 4) or the number of genetic variants (SNPs and small indels) across the whole genome (median 3 variants, IQR 1 to 8). Putative recombination events were detected in the bacterial genomes of 15% of individuals (n=117/791), accounting for 23% of the overall mutation count (n=1,721/7,577) and was predominantly located (n=1,367/1,721, 80%) in three prophages recombination hotspots within the reference genome used (NCTC8325)^30^ (Supplementary Figure 3).

### Genome-wide mutation enrichment analysis identifies evidence of adaptation

To identify loci in the *S. aureus* genome exhibiting evidence of parallel and convergent evolution that could represent potential signals of adaptation during colonisation, we applied a genome-wide mutation enrichment approach (Figure 1). Using clonal isolates sampled from the same host, we quantified the number of protein-altering mutations (missense, nonsense and frame-shift mutations) within each protein coding sequence (CDS) that arose *de novo* during *S. aureus* colonisation. We then statistically tested whether this was higher than expected when compared to the rest of the genome using a single-tailed Poisson test and correcting P values using a Benjamini & Hochberg correction for multiple testing.

**Figure 1.**
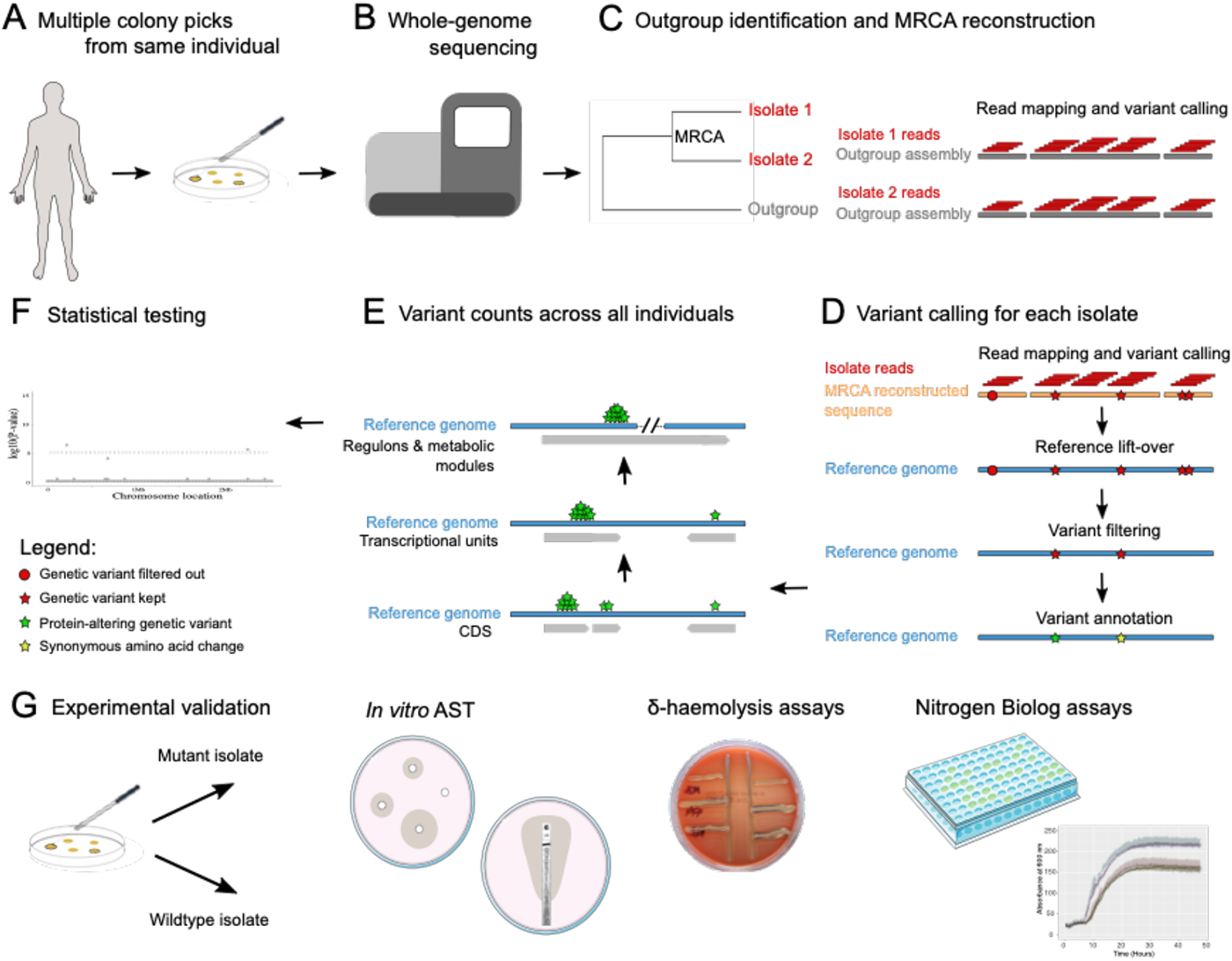
Design of genomic analyses to detect potential signals of adaptation. A. *S. aureus* colonies cultured from swabs taken from typical carriage sites of the same individual. B. Multiple isolates are whole genome sequenced from the same individual. C. A core-genome phylogeny is used to ensure isolates from the same host are clonal and to identify an appropriate outgroup. Isolate short reads are mapped to the outgroup assembly to call genetic variants. The sequence of the most recent common ancestor (MRCA) of all isolates from the same host is reconstructed. D. The short reads of each isolate are mapped to the MRCA reconstructed sequence to call variants wherein the reference allele represents the ancestral allele and the alternative allele the evolved one. The coordinates of variants in a complete and well-annotated reference genome (Reference lift-over) are determined. Variants on repetitive, low-complexity and phage regions are removed as well as those attributable to recombination (Variant filtering). In the last step, the effect of variants on genes is annotated (Variant annotation). E. The number of protein-altering mutations are counted on protein-coding genes (CDS), transcriptional units (operons) and high-level functional units across all individuals. F. Each functional unit is tested for an enrichment of protein-altering mutations compared to the rest of the genome. G. The mutant isolate (with a putative adaptive mutation) and a closely related wildtype isolate obtained from the same individual are tested *in vitro* for antibiotic susceptibility (AST), delta-haemolytic activity, and growth under a variety of nitrogen sources to validate the phenotypic effect of putative adaptive mutations.

Out of 2,326 CDS tested, only the genes encoding the accessory gene regulator A (*agrA*), the accessory gene regulator C (*agrC*) and the assimilatory nitrite reductase large subunit (*nasD*) showed a statistically significant (p-value <0.05 after adjusting for multiple testing) enrichment of protein-altering mutations (Figure 2A). Just below the genome-wide significance level, were genes encoding known antibiotic targets: *fusA* encoding the target of fusidic acid^31^; *dfrA* encoding the target of trimethoprim^32^; and *pbp2*, which encodes a target of beta-lactams^33^ (Figure 2A). The finding of *agr* genes (*agr*A and *agr*C), known to be frequently mutated in *S. aureus* carriers^34,35^, and that of known antibiotic targets demonstrated the feasibility of our approach in detecting putative adaptive mutations.

**Figure 2.**
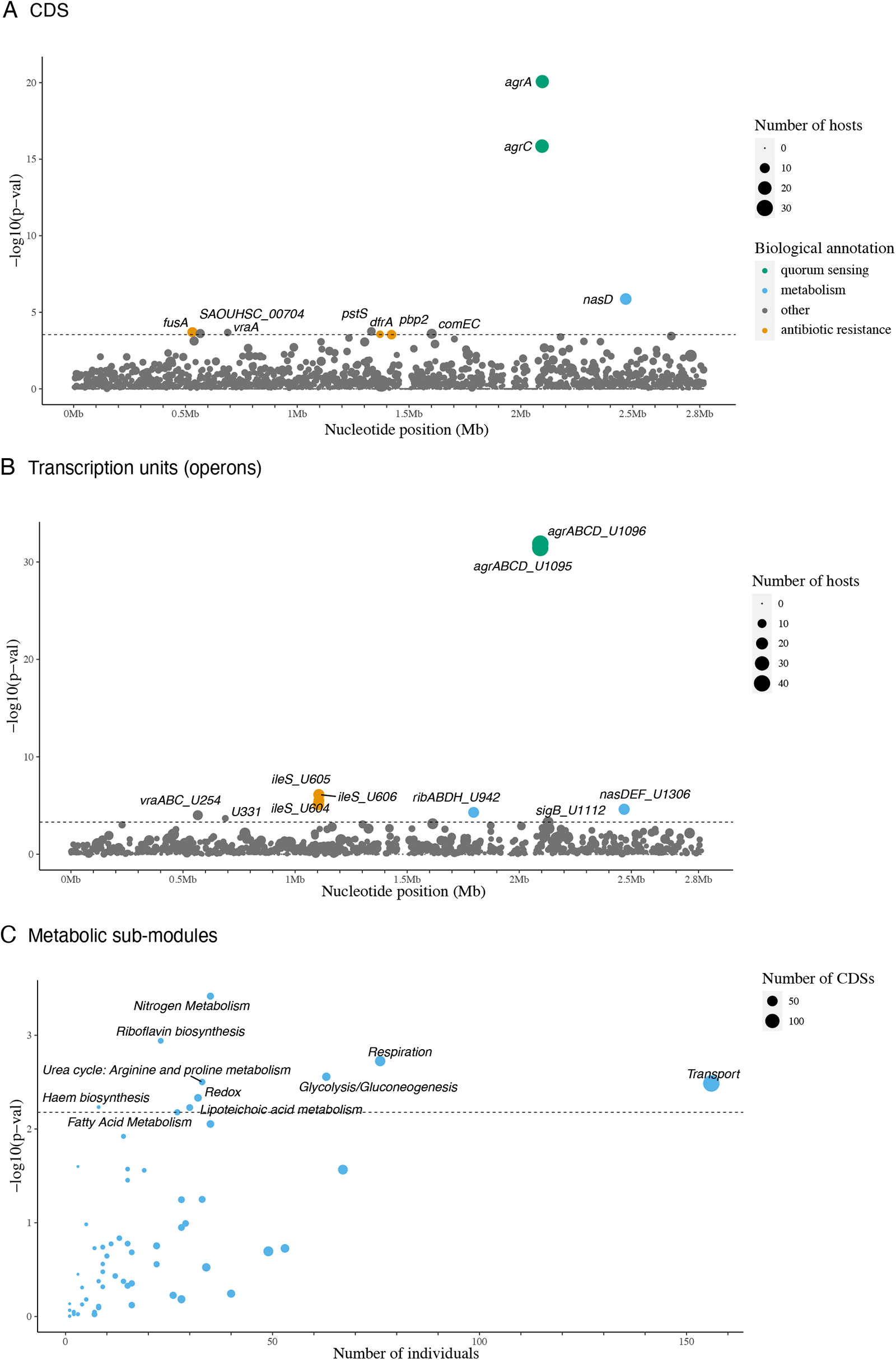
Loci enriched for protein-altering mutations in colonising isolates. (A) Protein coding sequences (CDS) and (B) transcriptional units (operons) enriched for protein-altering mutations in colonising isolates of the same host. Each circle denotes a single locus, whose size is proportional to the number of hosts mutations arose independently from. Loci are placed at the x-axis based on their chromosome coordinates, and at the y-axis based on their uncorrected p-value. The dotted horizontal line represents the genome-wide statistical significance threshold. (C) Metabolic sub-modules enriched for protein-altering mutations in colonising isolates. In the x-axis, number of independent acquisitions of protein-altering mutations in different hosts across all protein-coding sequences (CDS) of the same metabolic sub-module. The number of CDS making up metabolic sub-modules is indicated with the size of each circle. In the y-axis, strength of statistic association shown by adjusted p-value.

To broaden the search for signals of convergent evolution in groups of genes that are functionally related, we counted mutations among all genes belonging to the same transcription unit (operon)^36^. Out of 1,166 operons tested, nine reached statistical significance for an excess of protein-altering mutations (Figure 2B). These included three operons containing genes that reached statistically significance on their own: U1306 (*nasD*) and U1096 and U1095 (both containing *agrA* and *agrC*); and six additional operons containing CDS that did not reach statistical significance on their own: overlapping operons U605, U606 and U604, all containing the *ileS* gene; the U942 operon harbouring four riboflavin biosynthesis genes (*ribD*, *ribB*, *ribA* and *ribH*); the U254 operon containing genes involved in fatty acid metabolism (*vraA*, *vraB* and *vraC*); and the U331 operon which includes a single hypothetical protein (*SAOUHSC_00704*) (Supplementary Data 2).

At the highest functional level, we aggregated mutations within CDS of the same metabolic process, as defined by well-curated metabolic sub-modules in the *S. aureus* JE2 reference genome.^37^ Out of 61 metabolic pathways tested, 11 reached statistical significance for an excess of protein-altering mutations, with ‘nitrogen metabolism’ and ‘riboflavin biosynthesis’ pathways being the top two metabolic processes affected (Figure 2C), demonstrating a clear signal of selection on distinct metabolic processes.

### Nitrogen metabolic enzymes are enriched for mutations in colonising isolates

Nitrogen metabolism was the metabolic process most enriched by protein-altering mutations in colonisation isolates (Figure 2C). *nasD* (also named *nirB*) was the third most frequently mutated gene (in a total of 14 individuals), only after *agrA* (n=19) and *agrC* (n=20). *nasD* encodes the large subunit of the assimilatory nitrite reductase, an enzyme responsible for reducing nitrite (NO_2_-) to ammonium, an early step in the fixation of nitrogen from inorganic forms (Figure 3A). After *nasD*, the gene encoding the urease accessory protein UreG (*ureG*), was the second most mutated nitrogen metabolic enzyme (17^th^ hit, Supplementary Data 2). Urease is a nickel-dependent metalloenzyme that catalyses the hydrolysis of urea into ammonia (NH3) and carbon dioxide (CO_2_).^38^

**Figure 3.**
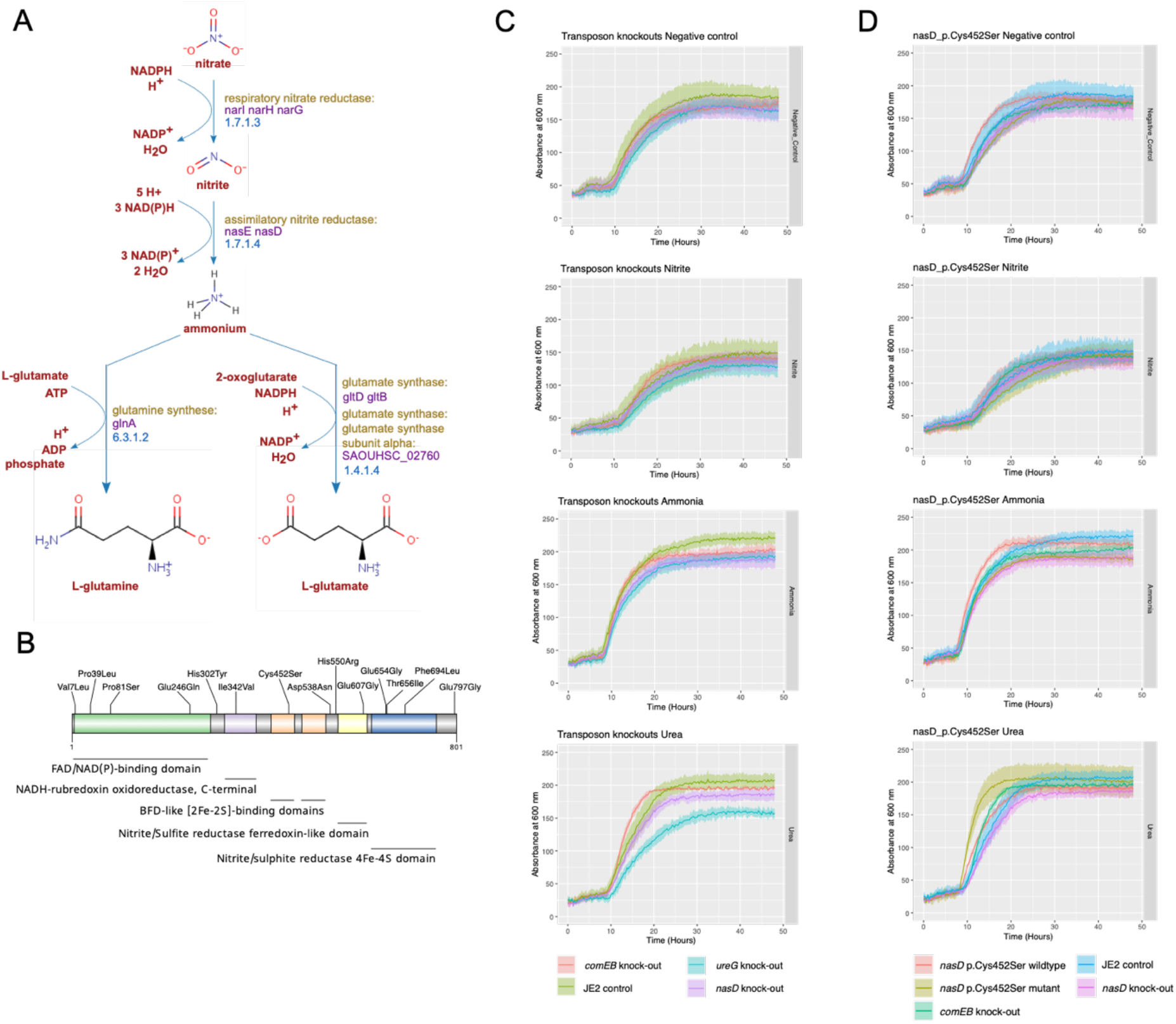
A. Role of the assimilatory nitrite reductase enzyme encoded by *nasD* in the nitrate assimilatory pathway of *S. aureus*. Adapted from BioCyc. B. Location of missense mutations along NasD protein. Pfam protein domains are shown in distinct colours. C. Growth curves of *S. aureus nasD/nirB* p.Cys452Ser mutant, wildtype (i.e. quasi-isogenic isolate lacking the *nasD/nirB* mutation from the same host), *nasD/nirB* knock-out, *comEB* knock-out (control) and JE (control) strain under the following nitrogen sources: negative control well, nitrite, ammonia, and urea. Coloured lines represent mean OD600 calculated across three replicates, and shaded coloured regions the standard deviation.

Because urea is by far the most abundant organic substance in nasal secretions^13^, we hypothesised that mutations in *nasD* and *ureG* could represent adaptations to the abundant availability of this nitrogen source. To investigate this, we first tested *nasD* and *ureG* transposon knockouts for their ability to grow under a variety of nitrogen sources. We observed rapid growth with amino acids like glycine, but slower growth with urea and ammonia as the primary nitrogen source, and even slower with nitrate and nitrite (Supplementary Figure 4, Supplementary Data 4). Compared to the control strain (*comEB* transposon knock-out), the growth of the *nasD* knock-out was compromised in multiple nitrogen sources (Figure 3C), including urea (growth rate 0.32 vs. 0.51, one-way ANOVA p-value < 0.01). Likewise, the growth rate of the *ureG* knock-out was significantly compromised with urea (0.25 vs. 0.51, one-way ANOVA p-value < 0.001, Supplementary Data 4), highlighting the critical role of *ureG* in the utilisation of urea as the main nitrogen source for growth.

Next, we tested available colonising isolates with naturally occurring *nasD* mutations (Supplementary Table 2), and their corresponding closely related *nasD-*wildtype isolates from the same host (n=3), for growth under the same nitrogen sources. Compared to the wildtype isolate, a Glu246Gln mutant (ST22) showed reduced growth under most nitrogen sources (Supplementary Figure 5), including in the negative control well, though the difference was most pronounced in urea, suggesting the fitness of this mutant was compromised relative to its wildtype. The Thr656Ile mutant (ST22) and wildtype both showed similar growth parameters across nitrogen sources, though the wildtype grew marginally better in urea than the mutant suggesting this mutation would be detrimental to growth in urea. Conversely, the Cys452Ser mutant (ST5) showed a statistically significant improvement in growth compared to its wildtype (in terms of a higher exponential growth rate: 0.64 vs 0.38, p-value < 0.001) in the presence of urea (Figure 3D), compared to inorganic nitrogen sources. These results point to an adaptive effect of *nasD* Cys452Ser mutation in the presence of urea. Interestingly, we also observed a strong effect of the strain’s genetic background on growth, with ST5 isolates (Supplementary Figure 5 I-L) growing comparably as well as the transposon control strains (ST8), and ST22 isolates growing comparably worse.

### Adaptive mutations reveal well-known and novel antibiotic resistance mutations

Our initial data suggested that the targets of antibiotics from distinct functional classes demonstrate potential signal of adaptation (Figure 2A). As such, we investigated whether mutations in these genes reduced susceptibility to their cognate antibiotics (Supplementary Figure 6) by testing the antibiotic susceptibility of closely related clinical isolates that were mutant and wild-type pairs from the same individual (Figure 1G). Mutations in *fusA* arose in 10 individuals. Out of the ten missense variants (Supplementary Table 3), five had the exact amino acid changes previously reported to confer fusidic acid resistance (Val90Ile, Val90Ala, Pro404Leu)^39^ or within the same codon (His457Arg) and were phenotypically resistant to fusidic acid. The other five isolates harbouring *fusA* missense variants were all susceptible to fusidic acid, ruling out an adaptive role of these mutations in fusidic acid resistance.

Five of the eight protein-altering mutations in *ileS* are known (Val588Phe and Val631Phe) or are in a codon (Gly593Ala) known to confer mupirocin resistance^39^ and exhibited elevated MICs compared to the wildtype clonal isolate from the same individual (Supplementary Table 3). We confirmed the role of a new frameshift mutation (Ile473fs) in mupirocin resistance (E-test MIC 1,024 μg/mL, breakpoint >12 μg/mL) and ruled out the effect of Gly591Ser (E-test MIC 0.5 μg/mL). Out of the five *S. aureus* isolates with missense variants in *dfrA,* three had amino acid changes reported to confer resistance to trimethoprim (His150Arg and two Phe99Tyr).^39^ The available isolate with Phe99Tyr was phenotypically resistant (MIC >=16 μg/mL), but the isolate carrying His150Arg was not (MIC <=0.5 μg/mL, zone diameter 27mm), ruling out the role of this mutation in trimethoprim resistance in this particular strain background.

Missense mutations in *pbp2* were all located within the transglycosylase domain of PBP2 (Supplementary Figure 6D), which is known to cooperate with PBP2A^40^ to mediate beta-lactam resistance in MRSA. The three PBP2-mutated strains from available collections^21^ were all ST22 (from phylogenetically distinct clades) MRSA (positive for *mecA*/PBP2a), but two were cefoxitin susceptible while retaining benzylpenicillin and oxacillin resistance (Supplementary Table 3). The corresponding PBP2-wildtype isolates from the same individual retain cefoxitin resistance, suggesting these mutations result in cefoxitin susceptibility.

We next investigated two sets of mutations putatively involved in glycopeptide resistance. First, *vraA,* a gene involved in fatty acid metabolism, was the seventh most mutated protein-coding gene (n=8 individuals), and is downregulated in daptomycin tolerant strains^41^. Mutations in other genes involved in cell membrane lipid metabolism (e.g., *mprF*/*fmtC* or *vraT*, Supplementary Table 4)^42^ are reported to reduce daptomycin susceptibility. Second, *pstS* a gene encoding a phosphate-binding protein, part of the ABC transporter complex PstSACB, was the fourth most frequently mutated protein-coding sequence (n=7 individuals) (Figure 2A). A point mutation in another phosphate transporter of *S. aureus* (*pitA*) increased daptomycin tolerance.^43^ We hypothesised that protein-altering mutations in *vraA* and *pstS* could have similar effect on daptomycin resistance. We determined daptomycin MICs and tolerance under a sub-inhibitory concentration of daptomycin (0.19 μg/mL) for the available *vraA*-mutated and *pstS*-mutated isolates (Supplementary Table 5), and with *pstS* and *vraA* loss-of-function (LOF) mutations from a larger collection (Supplementary Table 6). These results showed that neither the *pstS* or *vraA* mutations, or LOF mutations led to significant increases in daptomycin MIC, and only the mutant *pstS* p.Gln217* (mean AUC=10.5, one-way ANOVA p-value <0.01, Supplementary Data 3, Supplementary Figure 7) showed increased daptomycin tolerance. The absence of improved growth of mutants relative to controls indicates that the primary driver of *pstS* and *vraA* mutations was not daptomycin tolerance and suggests these mutations could be also metabolic adaptions to fatty acid metabolism.

### Agr-inactivating mutations arise frequently in colonising isolates

The genes encoding the sensor kinase AgrC and the response regulator AgrA were, by far, the most frequently mutated genes (Figure 2A), found in strains colonising 22 and 21 individuals, respectively (including one strain with both an ArgC and AgrA mutation). These genes belong to an operon encoding the accessory gene regulatory (Agr) system, a two-component quorum-sensing system that senses bacterial cell density and controls the expression of a number of important *S. aureus* virulence factors.^44^

In AgrC, protein-altering mutations were concentrated in the histidine kinase (HK) domain (n=16/20, Figure 4A), potentially abrogating phosphorylation of AgrA. For AgrA, mutations were enriched in the DNA binding domain (n=16/19, Figure 4B), likely preventing the binding of phosphorylated AgrA to its cognate DNA binding region. We additionally inspected mutations in the *agr* intergenic region and found that four of the five mutations in this region fall close to the AgrA binding site of Promoter 2 (Figure 4C). Altogether, these mutations likely abrogate expression of the Agr system by preventing the phosphorylation of AgrA or binding of phosphorylated AgrA to its cognate DNA binding region. To confirm this we tested putative *agr*-defective mutants, and their corresponding *agr*-wildtype isolate from the same host, for delta-haemolytic activity as a proxy for *agr* activity.^45^ Given the large number of mutations to test (Supplementary Table 7), we selected 24 isolates from available collections^21^ containing a representative mutation (i.e. missense, frameshift, stop gained and inframe indel) at each protein domain or intergenic region. As expected, the selected representative Agr-mutants were negative for delta-haemolytic activity, while their corresponding closely related wild-type isolates retained activity (Figure 4B).

**Figure 4.**
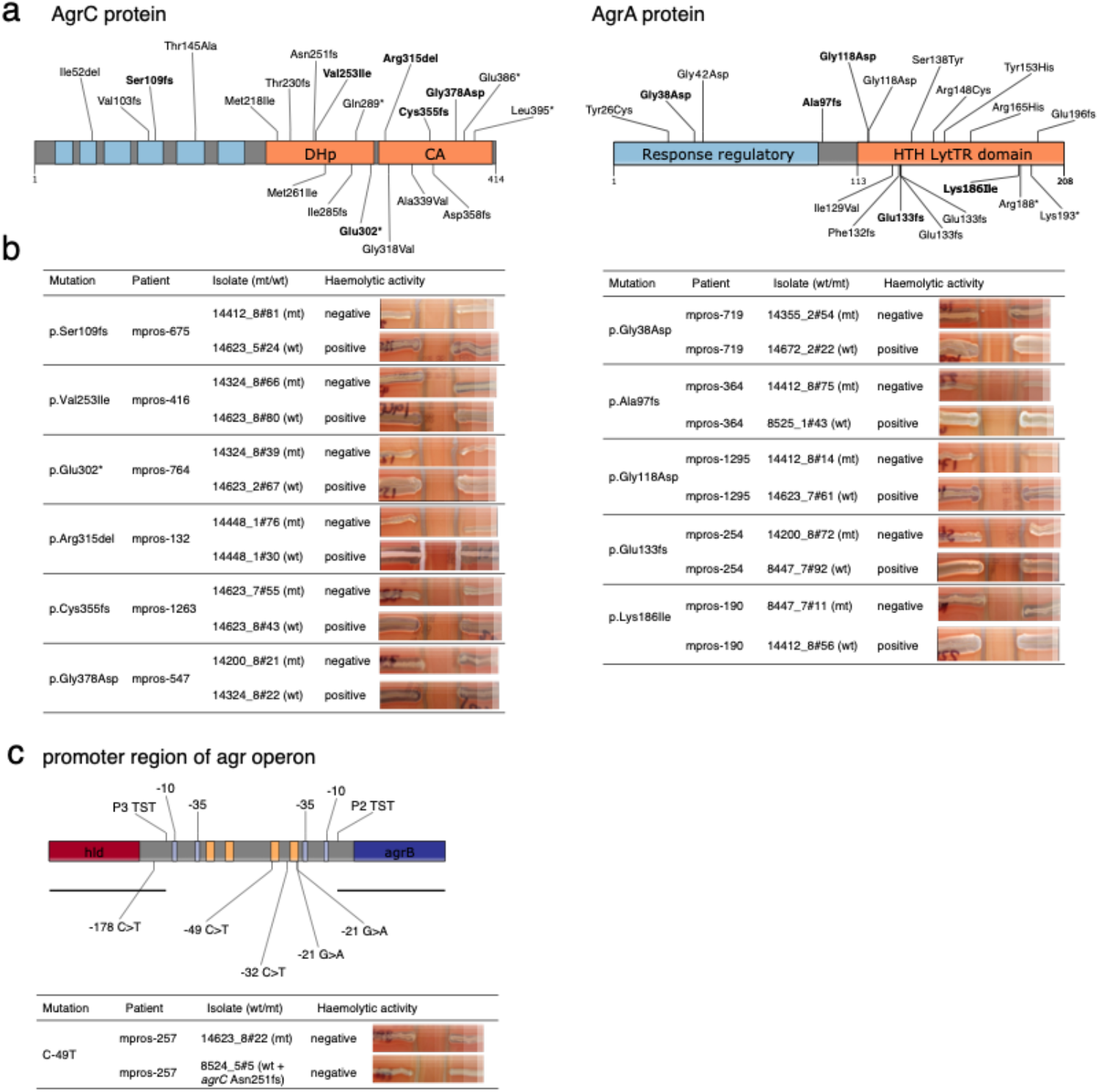
Mutations found in the accessory gene regulatory (Agr) system of colonizing strains. **(A)** Protein-altering mutations in the protein domains of the sensor kinase AgrC and the response regulator AgrA. The N-terminal sensor domain of AgrC comprises six transmembrane domains (coloured in blue) and is connected to a conserved C-terminal histidine kinase (HK) domain (coloured in orange). The HK domain is made up of two subdomains: the dimerization and histidine phosphotransfer (DHp) subdomain and the catalytic and ATP-binding (CA) subdomain.^79^ AgrA is comprised of a response regulatory domain (coloured in blue) and a DNA binding domain (coloured in orange). Isolates carrying mutations in bold were selected for haemolytic assays from available collections^21^ to represent different types of mutations (i.e. missense, frameshift, stop gained and inframe indel) at each protein domain. **(B)** Haemolytic activities of *S. aureus* isolates on sheep blood agar (SBA) plates used to test the activity of the Agr system. For each mutation, two isolates from the same host were tested, one carrying a selected Agr mutation (mutant) and a second isolate being wild type for the Agr system. A positive result is indicated by a widening of haemolysis seen in the region of RN4220. **(C)** Intergenic region containing agr promoters. The black horizonal lines represent the extent of transcript starting at the promoter 3 transcriptional start site (P3 TST), which encodes for RNAIII, and the transcript starting at promoter 2, which contains the whole *agrBDCA* coding region. Light blue boxes represent -10 and -35 boxes, whereas orange boxes the AgrA binding sites (“AgrA tandem repeats”). The only intergenic mutation carried by an available isolate (C-49T) yielded a negative haemolytic assay, as well as the isolate from the same host lacking this mutation, the latter attributable to a frameshift mutation in AgrC.

Mutations that inactivate *agr* have been reported in previous studies, predominantly in *agrC* and *agrA* genes, both in healthy carriers ^4,34,35^ and from multiple types of infections^45^, validating our approach to look for signals of adaptation. However, while some studies propose that agr-inactivating mutations arise more frequently in infected patients,^4^ others report similar frequencies in both infected and uninfected carriers.^35^ To investigate this, we tested whether *agr* mutants were more common in carriers who had staphylococcal infections compared to *S. aureus* uninfected carriers. We did not find this to be the case (p-value 0.17) after accounting for the number of sequenced isolates, genetic distance, collection, and clonal background as potential confounders (See Methods).

### Further putative adaptive mutations in an extended and larger dataset

Our original dataset was compiled in June 2019 (3,060 isolates from 791 individuals), to strengthen our initial findings, we searched for newly published studies with multiple colonisation isolates sequenced per individual (up to June 2023), to increase the sample size of the dataset and the chances of detecting novel adaptive variation. We applied the same curation, genomic and QC methodological steps to keep only high-quality and clonal genomes of the same individual from colonisation sources. A total of 4,090 additional isolate genomes obtained from 802 individuals and 15 different studies were included (Supplementary Table 8). Application of the genome-wide mutation enrichment approach to the combined dataset (7,150 isolates from 1,593 individuals) revealed even more genes reaching statistical significance for an excess of protein-altering mutations (Figure 5, Supplementary Figure 8), including the ones originally identified (*agrA*, *agrC* and *nasD*) plus an extra eight genes. The latter included *pstA* (which encodes for a nitrogen regulatory protein), *sasA* (*S. aureus* surface protein A also known as SraP (serine-rich adhesin for binding to platelets involved in adhesion and invasion)^46,47^ and *rsbU* (sigmaB regulation protein) and five genes yet to be functionally characterised (SAOUHSC_00704, SAOUHSC_00270, SAOUHSC_00621, SAOUHSC_02904 and SAOUHSC_00784). Genes encoding known antibiotic targets (*dfrA*, *fusA* and *pbp2*) remained among the top hits but below the genome-wide significance threshold. Among these was *mprF*, in which point mutations are known to confer daptomycin resistance. These results provide further evidence of the importance of nitrogen metabolism and identifies several uncharatersised genes likely to be critical for colonisation that warrant further experimental investigation.

**Figure 5.**
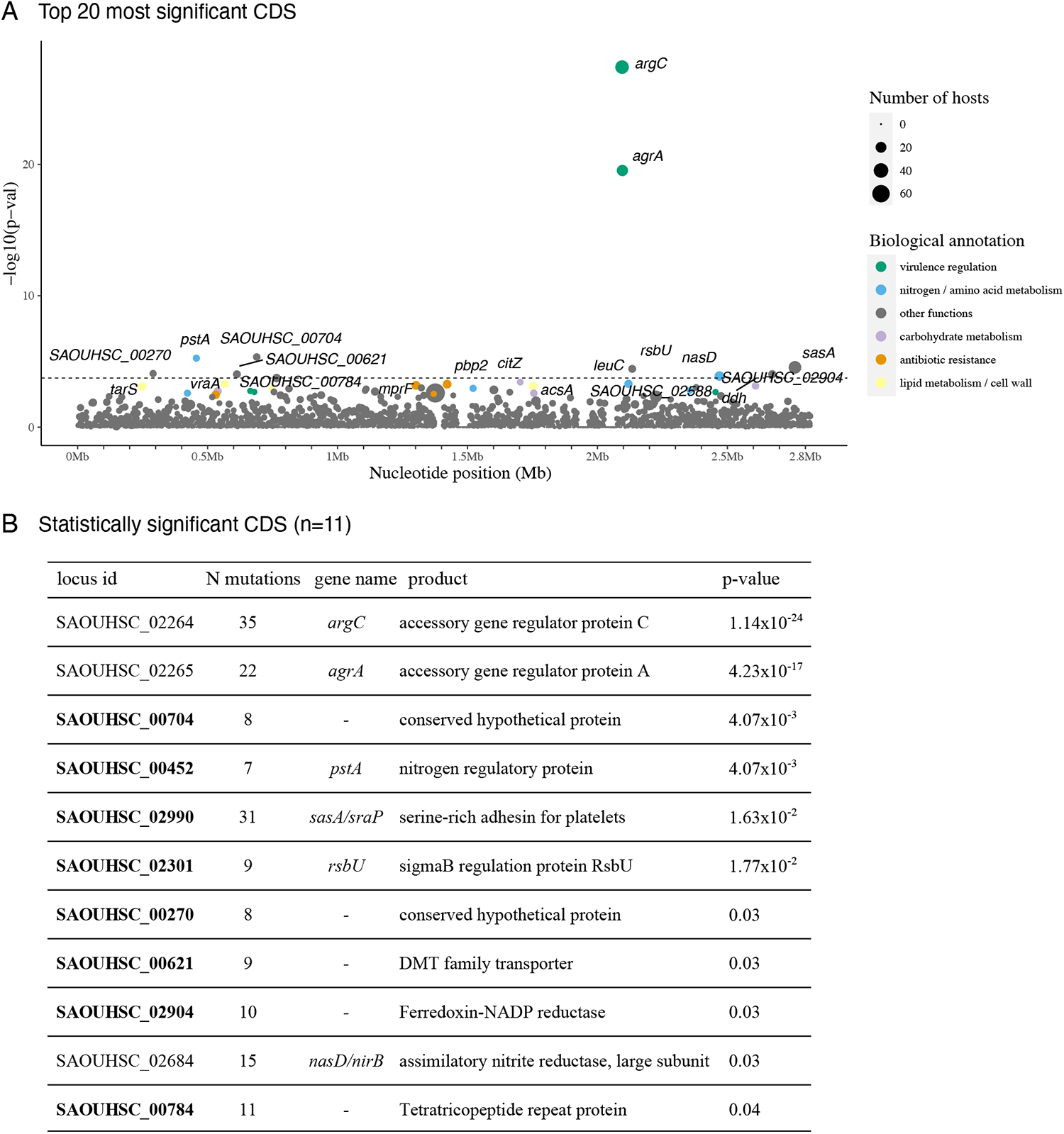
CDS enriched for protein-altering mutations in colonising isolates of the extended dataset. (A) The top 20 most significant CDS are labelled on the plot. Each circle denotes a single locus, whose size is proportional to the number of hosts mutations arose independently from. Loci are placed at the x-axis based on their chromosome coordinates, and at the y-axis based on their uncorrected p-value. The dotted horizontal line represents the genome-wide statistical significance threshold. (B) Locus id and annotation of statistically significant CDS (n=11) only. Locus ids in bold indicate the genes that became statistically significant in the extended dataset. The second column shows the number of mutations originating in different hosts. The p-value presented in this table corresponds to the Benjamini-Hochberg corrected p-value.

## Discussion

In this study we have provided a comprehensive view of the mutational landscape shaped by selective pressures that *S. aureus* is exposed to during human colonisation. The frequency and type of genomic mutations that arise provide a record of adaptive changes that commensal *S. aureus* underwent in response to evolutionary pressures in the host and provide novel insights into the biology of *S. aureus* in its primary niche. We compared the genomes of isolates collected from the same host, across a large number of hosts, to detect loci under parallel and convergent evolution.^48^

Our results provided indirect evidence of the ongoing metabolic adaptation of *S. aureus*, during colonisation, with the strongest selective pressure being on nitrogen metabolism. We observed that nitrogen metabolic enzymes are often mutated in colonising isolates, specifically genes encoding sub-units of the assimilatory nitrite reductase (*nasD*/*nirB*) and urease (*ureG*), and a nitrogen regulatory protein (*pstA*) in the extended dataset. Nitrite reduction can also be indicative of growth under anaerobic environments when nitrate (NO_3_-) and nitrite (NO_2_-) are used as terminal electron acceptors in place of O_2_.^49^ Indeed, genes related to dissimilatory nitrate and nitrite reduction are up-regulated under anaerobic conditions^50^, when *nasD*/*nirB* serves to detoxify the nitrite that accumulates in nitrate-respiring cells.^51^ Staphylococcal urease has also been implicated in adaptation to acid environments by ammonia production.^38^ Therefore, it cannot be ruled out for *nasD*/*nirB* and *ureG* mutations could represent adaptations to anaerobic and acidic environments, respectively.

The targets of fusidic acid (elongation factor G), trimethoprim (dihydrofolate reductase), mupirocin (isoleucyl-tRNA synthetase) and beta-lactams (penicillin-binding protein 2) showed a clear signal of adaptation as revealed by the independent emergence of mutations in the *S. aureus* isolates of multiple individuals. This most likely represents examples of directional selection, wherein *S. aureus* adapted to antimicrobial evolutionary pressures *in vivo*. This was supported by the identification of well-known resistance mutations in these genes, and concomitant reduced antibiotic susceptibility in isolates with these mutations, when compared to quasi ‘isogenic’ wild-type strains isolated from the same host. However, not all mutations detected in AMR loci were likely to be adaptive. This is exemplified by the characterisation of *fusA*, which had five mutations known to be involved in resistance leading to increases in MIC, and five never reported to cause resistance and not affecting fusidic acid susceptibility. It is therefore the excess of adaptive resistance-conferring mutations that increases the statistical significance of *fusA* and that of other AMR genes. We also identified novel mutations suggesting that the full diversity of resistance mutations to these drugs is yet to be fully understood and warrants further study. The mutations identified in the transglycosylase domain of PBP2, two of which resulted in cefoxitin susceptibility, are consistent with the cooperation of this native PBP with the acquired PBP2A to mediate beta-lactam resistance in MRSA, ^40^ and suggests that these might be compensatory mutations to optimise the function of the transglycosylase domain of PBP2.

Our results support previous observations that agr variation is selected for during colonisation.^35^ It has been proposed that a balance exists between wild-type and agr-defective cells in the population, where the latter, termed as ‘cheaters’, benefit from the secretions of wild-type cells without having to produce the costly cooperative secretions.^52^ However, in the context of colonisation, expression of the agr locus results in the down regulation of several surface proteins including cell wall secretory protein (IsaA)^53^ and fibronectin binding protein B (FnBPB)^54^ which are known to be involved in the attachment of *S. aureus* to cells in the nasal epithelium. Given the importance of these proteins to colonisation, it would be beneficial for *S. aureu*s populations to maintain subpopulations of cells that are primed for attachment should transmission to a new host occur. Thus, mutations in agr most likely represent an example of balancing selection, where the bacterial population as a whole benefits from having both active and defective agr systems, as opposed to a case of directional selection.

Our study has several limitations. First, the full genetic diversity of *S. aureus* in colonising sites was not captured by the datasets as we only had a median of two sequenced colonies available per individual. Having sequenced many more colonies, or directly from plate sweeps would have captured the full heterogeneity and provided a higher resolution picture of adaptation within bacterial sub-populations. Second, genetic changes identified between isolates of the same individual may not have arisen during colonisation of the sampled host (as assumed) but transmitted from another host, though these mutations still likely reflect recent diversification during colonisation. Third, we did not investigate changes in the gene content and large genetic re-arrangements, as those driven by movement of bacteriophages, between isolate genomes of the same host, this would require long-read sequencing. Fourth, we did not have metadata, such as antibiotic usage or the specific site of colonisation for 30.3% of isolates (e.g., multi-site screens). Finally, many of the isolates came from studies of *S. aureus* in hospital patients or with infections, which may have incorporated a bias towards mutations selected by antibiotics or other therapies. However, by increasing the overall sample size from ∼3,000 to ∼7,000 genomes we identified new genes significantly enriched for mutations including five currently uncharacterised genes and *sasA/sraP* which has not been previously been reported to be involved in colonisation, though it is known to mediate attachment to human cells.^46^ This suggests that studies using even larger sample sizes have the potential to identify further new signatures of adaption.

Future work focused on pre-defined patient groups (healthy colonised individuals), narrowly defined infection types^55,56^ with larger sample sizes and availability of host metadata will improve the identification of bacterial adaptive changes that promote survival in specific host niches and *in vivo* conditions; as well as pinning down strain/lineage-specific adaptations. Larger samples sizes will also allow us to determine which genes are essential for growth in different conditions, as shown by genes that are rarely inactivated.

While adaptation of clinical *S. aureus* strains during infection has been the focus of multiple recent studies,^4,20,57–59^ to our knowledge, this is the first comprehensive study to investigate adaptation of *S. aureus* populations experience during human colonisation. Our analysis has identified numerous metabolic pathways and genes likely critical to *S. aureus* colonisation that have not been previously reported to be involved in colonisation and demonstrated the functional impact of these mutations. Our data now warrant detailed experimental investigations to further elucidate *S. aureus* biology during colonisation. Finally, it is likely that our approach can be applied to other bacterial species with similar success.

## Methods

### Strain collections and data curation

We identified available collections of *S. aureus* genomes with multiple carriage isolates sequenced from the same human individual (Supplementary Data).^4,21–29^ The NCBI Short Read Archive (SRA) was systematically queried on June 2019 to identify BioProjects that met the following criteria (Figure 1): contained *S. aureus* genomic sequences, could be linked to a publication, included genomes of clinical isolates, clinical sources were known, multiple colonising isolates per host were sequenced, and host ids were available. Only isolates from colonisation specimens were kept, that is, from multi-site screens^21,25,27,28^ and typical colonising anatomical sites (nose^4,26,29^, armpit, groin, perineum and throat).^22–24^ Colonised hosts were classified as symptomatic or asymptomatic carriers based on whether they had a *S. aureus* infection or not, respectively. In studies where clinical specimens were systematically collected from recruited cases,^22,24,28^ individuals were labelled as asymptomatic carriers unless having a clinical specimen collected. In other studies, carriers were all explicitly referred to as infected^4^ or uninfected.^28,29^ In one study, only the nasal carriage controls were kept, as were thus labelled as uninfected. In the rest of studies, no information was available to determine their *S. aureus* infection status,^23,25,27^ and were thus labelled as ‘unknown’.

### Genomic analyses applied to all isolates

The Illumina short reads of all *S. aureus* genomes were validated using *fastqcheck* v1.1 (https://github.com/VertebrateResequencing/fastqcheck) and *de novo* assembled using Velvet v1.2.07^60^ to create draft assemblies. These were then corrected using the bacterial assembly and improvement pipeline^61^ to generate improved assemblies. QUAST v4.6.0^62^ was used to extract assembly quality metrics.

Sequence types (STs) were derived from improved assemblies by extracting all seven *S. aureus* multi-locus sequence type (MLST) loci and comparing them to the PubMLST database (www.PubMLST.org).^63^ Clonal complexes (CCs) were derived from these allelic profiles, allowing up to two allele mismatches from the reference ST. The short reads of each isolate were mapped to the same reference genomes (CC22 HO 5096 0412 strain, accession number HE681097) using *SMALT* v0.7.6 (http://www.sanger.ac.uk/resources/software/smalt/), whole-genome alignments were created by calling nucleotide alleles along the reference genome using *SAMtools* and *bcftools* v0.1.19.^64^ We kept the portion of the reference genome corresponding to the *S. aureus* core genome was kept in whole genome alignments to calculate core-genome pairwise SNP distances using *pairsnp* v0.0.1 (https://github.com/gtonkinhill/pairsnp). The core genome of *S. aureus*^65^ was derived from an independent, genetically and geographically diverse collection of 800 *S. aureus* isolates genomes from multiple host species^66^ using *Roary*^67^ with default settings. Core-genome alignments were used to construct a maximum likelihood phylogeny for each clonal complex using *RAxML* v8.2.8^68^ with 100 bootstraps.

### Genomic analyses applied to isolates of the same host

To avoid comparing the genomes of divergent strains from the same individual, only clonal isolates were kept for further analyses. Clonality was ruled out if isolates belonged to different clonal complexes or to the same clonal complex separated by more than 100 SNPs. Clonality was ruled in if isolates differed by less than the maximum within-host diversify previously reported (40 SNPs).^69^ Clonality was investigated for the remaining isolates pairs (differing between 40 to 100 SNPs) by making sure they all clustered within the same monophyletic clade in the phylogenetic tree.

The nucleotide sequence of the most recent common ancestor (MRCA) of all isolates of the same host was reconstructed first. To do this, we used the maximum likelihood phylogenies to identify, for each individual, the most closely related isolate sampled from a different individual that could be used as an outgroup. We used the *de novo* assembly of this outgroup isolate as a reference genome to map the short reads of each isolate, call genetic variants (SNPs and small indels) using *Snippy* v4.3.3 (https://github.com/tseemann/snippy), and build within-host phylogenies using *RAxML* phylogeny and rooted on the outgroup. The ancestral allele of all genetic variants at the internal node representing the MRCA of all isolates of the same host was reconstructed using PastML v1.9.20.^70^ This reconstructed ancestral sequence was used as the ultimate reference genome to call genetic variants (SNPs and small indels). This pipeline was implemented in four python scripts (identify_host_ancestral_isolate.step1.py to identify_host_ancestral_isolate.step4.py) available at https://github.com/francesccoll/staph-adaptive-mutations.

As variants were called in a different reference genome for each individual’s *S. aureus* strain, they had to be brought to the same reference genome to allow comparison and annotation across all individuals’ strains. We modified an already published script (*insert_variants.pl*)^48^ to find the genome coordinates of variants in the NCTC8325 (GenBank accession number NC_007795.1) and JE2 (NZ_CP020619.1) reference genomes. This script takes a 200-bp window around each variant in one reference (assembly) and finds the coordinates of this sequence in a new reference using *BLASTN*^71^ and *bcftools* v1.9^64^. Because of this requirement, variants at the edge of contigs (200 bp) were filtered out. The script was modified to keep the single best blast hit of each variant, meaning that variants with window sequences mapping to repetitive regions of the reference genome were removed. Variants in repetitive regions, detected by running Blastn v2.8.1+ on the reference genome against itself, and variants in regions of low complexity, as detected by *dustmasker* v1.0.0^72^ using default settings, were also filtered out. The final set of high-quality variants were annotated using *SnpEff* v4.3^73^ in both the NCTC8325 and JE2 reference genomes.

### Genome-wide mutation enrichment analysis

To scan for potential adaptive genetic changes recurrent across multiple individuals, we counted the number of functional mutations (i.e., those annotated as having HIGH or MODERATE annotation impact by *SnpEff*) in well-annotated functional loci across all individuals. Before that, putative recombination events, identified as variants clustered within a 1000-bp window in isolate genomes of the same host, were filtered out to avoid inflating mutation counts. When more than two isolates from the same host were available, we made sure the same mutations, identified in multiple case-control pairs of the same host, were counted only once.

We aggregated protein-altering mutations within different functional units. At the lowest level, we counted mutations within each protein coding sequence (CDS). To increase the power of detecting adaptive mutations in groups of genes that are functionally related, we aggregated mutations within transcription units (operons). The coordinates of transcription start and termination sites in the NCTC8325 reference genome were extracted from a study that comprehensively characterised the transcriptional response of *S. aureus* across a wide range of experimental conditions.^36,74^ To our knowledge, this is the best characterised reconstruction of transcriptional units in *S. aureus*. At the highest functional level, we aggregated mutations within CDS of the same metabolic process, as defined by well-curated metabolic sub-modules in the JE2 reference genome.^37^

We tested each functional unit (CDS, transcription unit and metabolic sub-module) for an excess of protein-altering (functional) mutations compared to the rest of the genome, considering the length of CDS, or cumulative length of CDS if testing high-order functional units involving multiple CDS. To do this, we performed a single-tailed Poisson test using the genome-wide mutation count per bp multiplied by the gene length as the expected number of mutations as previously implemented.^48^ Annotated features shorter than 300 bp long were not tested. P values were corrected for multiple testing using a Benjamini & Hochberg correction using the total number of functional units in the genome as the number of tests. We chose a significance level of 0.05 and reported hits with an adjusted P value below this value, unless otherwise stated.

### Other statistical analysis

We tested whether the presence of agr mutants, defined as isolates with protein-altering mutations in either *agrA* or *agrC*, was affected by hosts having an *S. aureus* infection (infection status). We fitted a binomial generalized linear model (GLM) using the presence of agr mutants as the binary response variable and *S. aureus* infection status as a binary predictor variable. We additionally included the number of sequenced isolates per host, genetic distance of these (expressed as the number of core-genome SNPs), collection and clonal background (clonal complex) as covariates to control for the effect of these potential confounders. This was implemented using the “glm” function (family binomial) in the base package within the statistical programming environment R version 3.4.1.^75^ The only predictors that increased the odds of detecting agr mutants were the number of sequenced isolates per host (odds ratio 1.20, 1.10 to 1.34 95% confidence interval, p-value < 0.001) and their genetic distance (odds ratio 1.06, 95% confidence interval 1.01 to 1.11, p-value < 0.05).

### In vitro antibiotic susceptibility testing

Isolates from frozen stocks were grown overnight on Columbia blood agar (CBA, Oxoid, UK) at 37°C. Fusidic acid or trimethoprim susceptibility testing was performed using disc (Oxoid, UK) diffusion as per EUCAST recommendations.^76^ Minimum inhibitory concentration (MIC) testing was performed for daptomycin, vancomycin and mupirocin. A loopful of the isolate added to phosphate buffered saline (PBS), adjusted to 0.5 McFarland, then a thin layer spread evenly on a Muller Hinton agar plate (Oxoid, UK). An antimicrobial gradient strip (Biomerieux, France) was carefully placed, then the plate incubated overnight at 37°C. The MIC was interpreted as the value on the strip above the point where growth stops.

### Biolog experiments

Isolates from frozen stocks were plated on to Lysogeny broth (LB) agar and grown overnight at 37°C. For the transposon knock-out strains, obtained from Nebraska transposon mutant library, the plates included 5ug/ml erythromycin. A damp swab was used to take sufficient colonies to create three 81% (+/-2%) turbidity solutions for each strain in 20ml PBS. 1.28ml of each turbid solution was added to 14.83 ml 1.2x IF0a (77268, Biolog), redox dye H (74228, Biolog), and PM3 Gram Positive Additive (made as described by the Biolog protocol). Each well of a PM3 plate (12121, Biolog) was inoculated with 150 µl of this solution. The inoculated plates were run on the Omnilog (Biolog) for 48 hours at 37°C. Readings were taken every 15 minutes.

### Delta-haemolysis experiments

The δ-haemolysis assay was performed as previously described.^45^ A thin streak of *Staphylococcus aureus* strain RN4220 was placed down the centre of a sheep blood agar plate. A thin streak of the test strain was placed horizontally up to, but not touching, RN4220. Test strains were tested in duplicate. Plates were incubated at 37°C for 18 hours, then at 4°C for 6 hours. Enhanced lysis by the test strain in the area near to RN4220 was an indicator of δ-haemolysis production.

### Growth curves

Test isolates were grown overnight at 37°C in tryptic soy broth (TSB) with 5ul/ml erythromycin (transposons) or TSB alone (non-transposons). The overnight cultures were then diluted 1/1000 in minimal media (1× M9 salts, 2 mM MgSO4, 0.1 mM CaCl2, 1% glucose, 1% casaminoacids, 1 mM thiamine hydrochloride and 0.05 mM nicotinamide) with 0.095ug/ml daptomycin. 300ul was added to a 96-well plate, then placed on a FluoStar Omega (BMG Labtech, Germany) for 24 hours incubation with shaking. Optical density measurement at OD_600_ was taken every 30 minutes, and standard curves produced. Each isolate was tested in biological and measurement triplicate.

The R scripts used to process raw growth data, plot growth curves, fit growth curves and compare growth parameters are available on GitHub (https://github.com/francesccoll/staph-adaptive-mutations/tree/main/growth_curves). Raw growth data (i.e. absorbance values at different time points) was processed with script prepare_growth_curves_data.R. Mean OD600 values and 95% confidence limits around the mean were plotted using ggplot2^77^ functions in script plot_growth_curves.R. Growth curves were fitted with Growthcurver^78^ and growth parameters (growth rate and area under the curve) extracted using script fit_and_plot_growth_curves.R. Due to the prolonged lag phase in curves obtained under daptomycin exposure, these curves were fitted after 7 hours. We fitted logistic curves to each replicate (n=9) using growthcurver package in R and extracted the growth rate and area under the logistic curve from fitted curves. These growth parameters were compared between isolates/strains (e.g. mutant vs. wildtype) using a one-way ANOVA to determine whether there were any statistically significant differences between the means (across replicates) of growth parameters between isolates (script: compare_growth_parameters.R).

## Supporting information

Manuscript

## Acknowledgements

This publication presents independent research supported by Wellcome grants 201344/Z/16/Z and 204928/Z/16/Z awarded to Francesc Coll. This publication was also supported by the Health Innovation Challenge Fund (WT098600, HICF-T5-342), a parallel funding partnership between the Department of Health and Wellcome Trust. EMH was supported by a UK Research and Innovation (UKRI) Fellowship: MR/S00291X/1. This publication was also supported by Wellcome Grant reference: 220540/Z/20/A, ‘Wellcome Sanger Institute Quinquennial Review 2021-2026’ and Wellcome Collaborative Award in Science: 211864/Z/18/Z. The views expressed in this publication are those of the author(s) and not necessarily those of the funders.

## Data availability statement

The whole genome sequences of the isolate collections used in this study are available on European Nucleotide Archive (ENA) under the accessions listed in Supplementary Data 1, which also includes isolate metadata. All scripts necessary to run the described analyses are available on GitHub (https://github.com/francesccoll/staph-adaptive-mutations). The full list of protein-coding regions, transcriptional units and metabolic processes enriched by protein-altering mutations can be found in Supplementary Data 2. Supplementary Data 3 and 4 include the data of bacterial growth curves.

## Author Contributions

Conceptualization: FC, EMH; Data curation: MT, FC; Formal bioinformatic analysis: FC, MM; Funding acquisition: FC, EMH, SJP; Investigation: FC, EMH; Bioinformatics methodology: FC, MM and DJ; Laboratory methodology: BB, KB and EMH; Project administration: EMH and SJP; Resources: JP, EMH and SJP; Supervision: EMH, JAG, JP and SJP; Validation: BB, KB, RCM; Visualization: FC; Writing – original draft: FC and EMH; Writing – review & editing: all authors.

## Notes

### Competing Interest Statement

The authors have declared no competing interest.

