## Supplementary material for "The mutational landscape of *Staphylococcus aureus* during colonisation": Manuscript

### The mutational landscape of *Staphylococcus aureus* in its primary niche

#### Supplementary Tables

Supplementary Table 1. Collections with multiple sequenced colonising isolates available from the same individual used in this study

Supplementary Table 2. Protein-altering mutations in *nasD* (*nirB*) identified in colonising isolates of the same host

Supplementary Table 3. Putative adaptive mutations in genes encoding for antibiotic targets identified in colonising isolates of the same host

Supplementary Table 4. Reported daptomycin-resistant mutations in *S. aureus*

Supplementary Table 5. Hypothesised daptomycin adaptive mutations in colonising isolates of the same host

Supplementary Table 6. Loss-of-function mutations in *pstS* and *vraA* genes found in a collection of 2,345 MRSA isolates

Supplementary Table 7. Protein-altering mutations in the AgrCA two-component system identified in colonising isolates of the same host

Supplementary Table 8. Collections with multiple sequenced colonising isolates available from the same individual identified after June 2019

**Supplementary Figures**

Supplementary Figure 1. Selection criteria used to identify collections of multiple colonising isolates sequenced per host

Supplementary Figure 2. Genetic diversity between colonising isolates of the same host

Supplementary Figure 3. Density of mutations attributable to recombination

Supplementary Figure 4. Growth curves of *S. aureus nasD* and *ureG* knock-out mutants under different nitrogen sources

Supplementary Figure 5. Growth curves of *S. aureus nasD* (*nirB)* mutants under different nitrogen sources

Supplementary Figure 6. Protein-altering mutations detected in known antibiotic targets.

Supplementary Figure 7. Growth curves of *pstS* and *vraA* mutant and wildtype S. aureus clinical isolates under daptomycin exposure

Supplementary Figure 8. Loci enriched for protein-altering mutations in the extended dataset

**Supplementary files:**

Supplementary Data 1. Isolate accession and metadata.

Supplementary Data 2. Hits of mutation enrichment analyses.

Supplementary Data 3. Raw growth curves measurements and growth parameters obtained with and without daptomycin

Supplementary Data 4. Raw growth curves measurements and growth parameters obtained under different nitrogen sources

Supplementary Table 1. Collections with multiple sequenced colonising isolates available from the same individual used in this study

| Study Accession (Publication) | Isolates used (out of available) | Isolates kept after QC | Sources of Isolation | Years of Isolation | Place of Isolation | Number of hosts | Median isolates per individual |
|---|---|---|---|---|---|---|---|
| PRJEB3174[1] | 727/2,282 | 702 | Multi-site screen | 2012 - 2013 | East of England, UK | 284 | 2 (IQR 2-3) |
| PRJNA324190[2] | 1,338/1,977 | 1,107 | Nose, perineum, groin, throat and armpit | 2011 - 2012 | Brighton, UK | 261 | 3 (IQR 2-6) |
| PRJNA369475[3] | 566/1,163 | 551 | Nasal swabs | 2009 - 2013 | Oxford and Brighton, UK | 105 | 5 (IQR 5-5) |
| PRJEB9390[4] | 172/383 | 139 | Groin, nose, axilla | 2014 | Singapore | 78 | 2 (IQR 2-2) |
| PRJEB2862[5] | 159/275 | 132 | Nose, perineum and groin | 2010 - 2011 | Brighton, UK | 67 | 2 (IQR 2-2) |
| ERP000130[6] | 143/172 | 96 | Multi-site screen | 2008 | Thailand | 22 | 2 (IQR 2-2) |
| PRJEB20148[7] | 90/262 | 89 | Nasal swabs | 2015 | UK | 18 | 5 (IQR 5-5) |
| PRJEB11177[8] | 35/448 | 22 | Nose, throat and perineum | 2011 - 2012 | London, UK | 17 | 2 (IQR 2-2) |
| PRJEB2655 and PRJEB7654[9] | 97/165 | 89 | Multi-site and nasal swabs | 2012 - 2014 | Galway, Ireland and Cambridge, UK | 16 | 3 (IQR 2-5) |
| PRJEB4141[10] | 170/172 | 133 | Nasal swabs | Not found | England | 4 | 52 (IQR 39-55) |
| Total | 3,497/7,299 | 3,060 | | | | 872 | 2 (IQR 2-4) |

Collections of *S. aureus* genomes used in this study where multiple isolate genomes were available from the same host. Only isolate genomes kept after quality control were used for further analyses.

Supplementary Table 2. Protein-altering mutations in *nasD* (*nirB*) identified in colonising isolates of the same host

| Gene (locus tag) | Chr. position | Nucleotide change | Annotation | Amino acid change | Protein domain* | Patient id (isolate id) | Selected for testing** |
|---|---|---|---|---|---|---|---|
| *nasD*/*nirB* (SAOUHSC_02684) | 2469191 | T > C | missense | p.Glu797Gly | - | young2017-P085 (SRR5250431) | not available |
| | 2469499 | G > T | missense | p.Phe694Leu | domain E | price2016-2058578 (SRR3729258) | not available |
| | 2469614 | G > A | missense | p.Thr656Ile | domain E | mpros-381 (8525_1#72) | selected |
| | 2469620 | T > C | missense | p.Glu654Gly | domain E | young2017-P095 (SRR5250387) | not available |
| | 2469761 | T > C | missense | p.Glu607Gly | domain D | price2016-H240 (SRR3728572) | not available |
| | 2469932 | T > C | missense | p.His550Arg | - | price2016-3424520 (SRR3728593) | not available |
| | 2469969 | C > T | missense | p.Asp538Asn | - | harrison2016-GR11 (13414_7#15) | not available |
| | 2470227 | A > T | missense | p.Cys452Ser | domain C | mpros-194 (9716_3#67) | selected |
| | 2470557 | T > C | missense | p.Ile342Val | domain B | price2016-H149 (SRR3728524) | not available |
| | 2470677 | G > A | missense | p.His302Tyr | - | price2016-3414160 (SRR3730410) | not available |
| | 2470845 | C > G | missense | p.Glu246Gln | domain A | mpros-547 (14324_8#22) | selected |
| | 2471340 | G > A | missense | p.Pro81Ser | domain A | price2016-H127 (SRR3728586) | not available |
| | 2471465 | G > A | missense | p.Pro39Leu | domain A | price2016-H179 (SRR3729445) | not available |
| | 2471562 | C > A | missense | p.Val7Leu | domain A | chow2017-CD140187 (17059_1#16) | not available |

*Domain referred here: domain A, FAD/NAD(P)-binding domain; domain B, NADH-rubredoxin oxidoreductase domain; domain C, BFD-like [2Fe-2S]-binding domains; domain D, Nitrite/Sulfite reductase ferredoxin-like domain; domain E, Nitrite/sulphite reductase 4Fe-4S domain. **Only isolates from internal collections (i.e. coll2017) were available for testing.

Supplementary Table 3. Putative adaptive mutations in genes encoding for antibiotic targets identified in colonising isolates of the same host

| Antibiotic | Gene | Chr. position | Nt. change | Amino acid change | Patient id (isolate id) | Known role* | Phenotypic resistance (MIC)** | Wildtype isolate id | Wildtype isolate MIC ** |
|---|---|---|---|---|---|---|---|---|---|
| Fusidic acid | *fusA* | 531299 | G > A | p.Val90Ile | coll2017-1295 (14623_7#61) | Yes | R (2 µg/mL) | 14412_8#14 | S (<=0.5 µg/mL) |
| | | 531300 | T > C | p.Val90Ala | young2017-P040 (SRR5249845) | Yes | Isolate NA | - | - |
| | | 531392 | C > T | p.Arg121Cys | coll2017-608 (14412_8#87) | No | S (<=0.5 µg/mL) | 14355_1#58 | S (<=0.5 µg/mL) |
| | | 531719 | C > T | p.Leu230Phe | coll2017-534 (14672_2#27) | No | S (<=0.5 µg/mL) | 14200_6#68 | S (<=0.5 µg/mL) |
| | | 532242 | C > T | p.Pro404Leu | coll2017-254 (8447_7#92) | Yes | R (16 µg/mL) | 14200_8#72 | S (<=0.5 µg/mL) |
| | | 532401 | A > G | p.His457Arg | coll2017-1162 (14208_8#9) | Codon | R (>=32 µg/mL) | 14623_3#77 | R (16 µg/mL) |
| | | 532401 | A > G | p.His457Arg | coll2017-527 (14208_8#16) | Codon | R (>=32 µg/mL) | 14200_6#27 | R (16 µg/mL) |
| | | 532550 | G > A | p.Gly507Ser | coll2017-104 (14448_1#44) | No | S (<=0.5 µg/mL) | 14200_8#74 | S (<=0.5 µg/mL) |
| | | 532770 | C > T | p.Ala580Val | coll2017-103 (14355_1#61) | No | S (<=0.5 µg/mL) | 8524_3#66 | S (<=0.5 µg/mL) |
| | | 533048 | G > A | p.Asp673Asn | young2017-P094 (SRR5249970) | No | Isolate NA | - | - |
| Mupirocin | *ileS* | 1107196 | A > G | p.Lys23Glu | price2016-H277 (SRR3728914) | No | Isolate NA | - | - |
| | | 1108541 | DEL.[2] | p.Ile473fs | coll2017-121 (14324_8#70) | No | R (1,024 µg/mL)§ | 8490_3#36 | S (0.19 µg/mL)§ |
| | | 1108891 | G > T | p.Val588Phe | coll2017-125 (14623_6#41) | Yes | R (>=512 µg/mL) | 14200_8#87 | S (<=2 µg/mL) |
| | | 1108891 | G > T | p.Val588Phe | coll2017-291 (14208_8#53) | Yes | I (12 µg/mL) § | 8524_5#51 | S (0.19 µg/mL)§ |
| | | 1108900 | G > A | p.Gly591Ser | coll2017-520 (14355_1#12) | No | S (0.5 µg/mL) § | 14200_6#18 | S (0.19 µg/mL)§ |
| | | 1108907 | G > C | p.Gly593Ala | price2013-X035 (SRR2053963) | Codon | Isolate NA | - | - |

| | | | | | | | | |
|---|---|---|---|---|---|---|---|---|
| | | 1109020 | G > T | p.Val631Phe | coll2017-177 (14623_8#7) | Yes | I (3 µg/mL)§ | 14324_8#86 | S (0.19 µg/mL)§ |
| | | 1109020 | G > T | p.Val631Phe | coll2017-75 (14623_8#86) | Yes | I (12 µg/mL)§ | - | - |
| Trimethoprim | *dfrA* | 1370041 | T > C | p.His150Arg | coll2017-789 (14672_2#49) | Yes | S (<=0.5 µg/mL, 27 mm) | 14200_7#34 | S (<=0.5 µg/mL, 28 mm) |
| | | 1370098 | T > C | p.Asp131Gly | price2016-H213 (SRR3730594) | No | Isolate NA | - | - |
| | | 1370180 | C > A | p.Asp104Tyr | chow2017-CD140174 (17059_1#8) | No | Isolate NA | - | - |
| | | 1370194 | A > T | p.Phe99Tyr | price2016-H264 (SRR3730705) | Yes | Isolate NA | - | - |
| | | 1370194 | A > T | p.Phe99Tyr | coll2017-237 (14623_6#13) | Yes | R (>=16 µg/mL) | 8447_7#65 | S (<=0.5 µg/mL) |
| Beta-lactams | *pbp2* | 1421571 | C > A | p.Ser7Tyr | price2016-2107188 (SRR3728791)[3] | - | Isolate NA | - | - |
| | | 1421757 | T > C | p.Leu69Ser | chow2017-CD140955 (16870_4#6)[4] | - | Isolate NA | - | - |
| | | 1421900 | C > T | p.Arg117Cys | young2017-P069 (SRR5251343)[3] | - | Isolate NA | - | - |
| | | 1421900 | C > A | p.Arg117Ser | coll2017-292 (8524_5#53)[4] | - | CFX R (>4 µg/mL) | 14623_2#62 | CFX R (>4 µg/mL) |
| | | 1421933 | C > A | p.Arg128Ser | young2017-P069 (SRR5250494)[3] | - | Isolate NA | - | - |
| | | 1421972 | T > G | p.Phe141Val | coll2017-547 (14324_8#22)[4] | - | CFX S (<=4 µg/mL) | 14200_8#21 | CFX R (>4 µg/mL) |
| | | 1421976 | G > A | p.Gly142Asp | young2017-P080 (SRR5250797)[3] | - | Isolate NA | - | - |
| | | 1421994 | C > T | p.Thr148Ile | young2017-P079 (SRR5250247)[3] | - | Isolate NA | - | - |
| | | 1422237 | G > C | p.Gly229Ala | young2017-P006 (SRR5249848)[3] | - | Isolate NA | - | - |
| | | 1422256 | C > A | p.Asn235Lys | coll2017-150 (8490_3#60)[4] | - | CFX S (<=4 µg/mL) | 14448_3#45 | CFX R (>4 µg/mL) |

* Known antibiotic-resistant mutations were extracted from Kumar *et al.* 2020[11].

** Isolates from 'coll2017-' patients (internal collection) had available AST VITEK data and were available for further antibiotic susceptibility testing. A few isolates were re-tested with disc diffusion, for which zone diameters are indicated in millimetres. §MIC confirmed using E-test. Isolate NA: isolate not available for re-testing.
[2] Deletion: GCGAAATTATCATGA > G. [3] MSSA strain (lacking both *mecA* and *mecC*). [4] MRSA strain with *mecA*. Abbreviations: MIC, Minimum Inhibitory Concentration; CFX, cefoxitin; AST, antibiotic susceptibility testing.

Supplementary Table 4. Reported daptomycin-resistant mutations in *S. aureus*

| Gene | Mutations reported in the literature | Locus tag (hit position) | Mutations in within-host dataset |
|---|---|---|---|
| *mprF* | T345A[12–17], P314L[13,18,19], T345I[13,14,18,20–22], T345X, V351E[23], S295L[19,22,24,25], L338S[26], S337L[14,18,19,22,26–28], L776S[14], A475P[14], L459_H466 del[14], L291I[14], W424R[14], L341S[14], P314L[19], S295A[27], I348del[27], R50L[18], R301L[18], L425F[18], P314L[29], L826F[14,19,22,29–31], L826I[19], S829L[32], H376Y&W424C[19], A302V[19], M347R[19], 41insN[19], M347L[33], L770F[33], I420N[22], G61V[22]<br>Increased expression[16,33] | SAOUHSC_01359<br>19th hit | p.Arg50Cys, p.Gly61Val, p.Gly61Val, p.Ser156Tyr, p.Met323Ile, p.Glu593Asp, p.Ala715Glu, p.Ala727Thr, p.Ser829Leu, p.Ser829Leu |
| *yycF* | K151N[19] | SAOUHSC_00020<br>Not detected | Not extracted |
| *yycG* | S221P[13], R263C[13,15,17], I568V[26], I185T[26], Q51K[26], G199E[19], 369delQ[19], G223D[34], M426I[31]<br>Increased expression[35] | SAOUHSC_00021<br>354th | Not extracted |
| *rpoB* | I953S[13], A1086V[13,15,17] L770F[33],Q468K[36], A477D[25,36], H481N[36], H481Y[16,23], H481R[16], S464P[36], A621E[16,37] R484C&N641K[34], D471Y&A473S&A477S&E478D[34] | SAOUHSC_00524<br>1421st | Not extracted |
| *rpoC* | F632S[13], Q961K[13,15,17], N735K[19] | SAOUHSC_00525 | Not extracted |

| | | | |
|---|---|---|---|
| | | 1068th | |
| *vraS* | Increased expression[25,26,30,38,39]<br>T331I[32], G45V[38], E276K[40], L114S&D242G[41] | SAOUHSC_02099<br>2035th | Not extracted |
| *vraR* | Increased expression[25,30,33,39] | SAOUHSC_02098<br>2034th | Not extracted |
| *vraT* | A151T[42] | SAOUHSC_02100<br>1007th | Not extracted |
| *tagH* | A39T[42] | SAOUHSC_02009<br>1633rd | Not extracted |
| *cls2* | R320L[19], R320S[19], F85X[19], L77F[19], A56G&T58N[19], A23V[22,43], T33N[43], L52F[22,43], F60[22,43], E38G[21], L190F[21]<br>Increased expression[44] | SAOUHSC_02323<br>915th | Not extracted |
| *pgsA* | A64V[22,45], S177F[22,45], K65R[22], V59N[22,45], V59D[45], G61S[45], K75N[45], K135E[31,45], D187E[45] | SAOUHSC_01260<br>327th | Not extracted |
| *ddl* | No mutations found | SAOUHSC_02318<br>1252nd | Not extracted |
| *dltA* | A426E[19]<br>Increased expression[16,44,46] | SAOUHSC_00869<br>49th | p.Ser37Ala<br>p.Leu219Ile<br>p.Asp306Tyr<br>p.Thr310Ile<br>p.Pro467Ser<br>p.Val482Ala |
| *clpX* | A348V[21], nt168_del[25] | SAOUHSC_01778<br>177th | Not extracted |

Mutations reported to be associated be daptomycin non-susceptibility in the literature. The hit position of candidate genes in the CDS mutation enrichment analysis is presented in the third column, along with the mutations found in this dataset in the fourth column.

Supplementary Table 5. Hypothesised daptomycin adaptive mutations in colonising isolates of the same host

| Gene (locus tag) | Chr. position | Nucleotide change | Annotation | Amino acid change | Protein domain* | Patient id (isolate id) | Selected for testing | Dap. MIC | Daptomycin MIC of wildtype isolate** |
|---|---|---|---|---|---|---|---|---|---|
| *pstS* (SAOUHSC_01389) | 1331476 | C > T | missense | p.Ala291Thr | - | price2016-2754372 (SRR3731418) | NA | NA | NA |
| | 1331536 | A > G | missense | p.Phe271Leu | domain A | price2016-H123 (SRR3731460) | NA | NA | NA |
| | 1331730 | C > T | missense | p.Gly206Glu | domain A | mpros-359 (8525_1#49) | Yes | 0.38 | 0.5 (14208_8#2) |
| | 1331764 | C > T | missense | p.Ala195Thr | domain A | young2017-P048 (SRR5250629) | NA | NA | NA |
| | 1331852 | TGGTG > T | frameshift | p.Ser164fs | domain A | mpros-64 (14200_8#1) | Yes | 0.38 | 0.38 (8524_3#24) |
| | 1331853 | G > T | missense | p.Pro165Gln | domain A | price2016-H187 (SRR3729045) | NA | NA | NA |
| | 1332186 | T > C | missense | p.Glu54Gly | domain A | young2017-P046 (SRR5250023) | NA | NA | NA |
| *vraA* (SAOUHSC_00557) | 566249 | C > T | missense | p.Pro60Ser | domain A | mpros-97 (8524_3#62) | Yes | 0.19 | 0.5 (14448_1#41) |
| | 566255 | C > G | missense | p.Gln62Glu | domain A | chow2017-CD140187 (17059_1#17) | NA | NA | NA |
| | 566441 | C > A | missense | p.His124Asn | domain A | price2016-H122 (SRR3729370) | NA | NA | NA |
| | 566794 | A > AT | frameshift | p.Ser244fs | domain A | young2017-P009 (SRR5250318) | NA | NA | NA |
| | 567014 | A > G | missense | p.Ile315Val | domain A | harkins2018-SS_157 (ERR1904049) | NA | NA | NA |
| | 567321 | T > A | stop gained | p.Leu417* | domain B | mpros-381 (14208_8#20) | Yes | 0.75 | 0.38 (8525_1#72) |
| | 567321 | TA > T | frameshift | p.Lys419fs | domain B | paterson2015-Staff_D (10770_3#44) | Yes | 0.75 | 0.38 (10900_1#28) |
| | 567390 | C > A | missense | p.Ala440Glu | domain B | chow2017-CD140764 (16870_3#11) | NA | NA | NA |

*pstS contains a single PBP superfamily domain (http://pfam.xfam.org/protein/Q2FYP6), here labelled as domain A. vraA is made up of two domains (http://pfam.xfam.org/protein/Q2G0K3): an AMP-binding enzyme domain (here labelled as domain A) and an AMP-binding enzyme C-terminal domain (domain B). ** isolates from the same host without mutation.

Supplementary Table 6. Loss-of-function mutations in *pstS* and *vraA* genes found in a collection of 2,345 MRSA isolates

| Gene (locus tag) | Chr. position | Nuc. change | Annotation | Amino acid change[1] | Protein domain[2] | isolate id | Dap. MIC[3] | Related isolate | Genetic distance (SNPs) | Related isolate Dap. MIC[3] | Known dap. mutations |
|---|---|---|---|---|---|---|---|---|---|---|---|
| *pstS* (SAOUHSC_01389) | 1387079 | A > AT | frameshift | p.Ter328fs | - | 14355_1#80 | ND | 14623_5#49 | 73 | ND | - |
| | 1387079 | A > AT | frameshift | p.Ter328fs | - | 14623_7#5 | ND | 14623_5#49 | 83 | ND | - |
| | 1387339 | GT > G | frameshift | **p.Thr241fs** | domain A | 14324_8#29 | 0.5 | 14448_2#11 | 16 | 0.5 | - |
| | 1387339 | G > GT | frameshift | **p.Thr241fs** | domain A | 14623_5#47 | ND | 14355_2#16 | 86 | ND | - |
| | 1387339 | GT > G | frameshift | **p.Thr241fs** | domain A | 14672_2#74 | ND | 14623_3#79 | 53 | ND | - |
| | 1387339 | GT > G | frameshift | **p.Thr241fs** | domain A | 8447_7#72 | ND | 14623_3#79 | 52 | ND | - |
| | 1387412 | G > A | stop gained | p.Gln217* | domain A | 14412_8#80 | 0.5 | 14355_1#89 | 16 | 0.5 | - |
| | 1387491 | A > AT | frameshift | **p.Asn190fs** | domain A | 14355_1#57 | ND | 14623_2#41 | 72 | ND | - |
| | 1387491 | A > AT | frameshift | **p.Asn190fs** | domain A | 14355_2#77 | ND | 14623_2#41 | 74 | ND | - |
| | 1387491 | AT > A | frameshift | **p.Asn190fs** | domain A | 14448_3#24 | 0.25 | 14200_6#37 | 42 | 0.5 | - |
| | 1387491 | A > AT | frameshift | **p.Asn190fs** | domain A | 14555_7#38 | ND | 14623_2#41 | 74 | ND | - |
| | 1387491 | AT > A | frameshift | **p.Asn190fs** | domain A | 14623_7#4 | 0.38 | 14200_6#37 | 41 | 0.5 | - |
| | 1387566 | TGGTG > T | frameshift | p.Ser164fs | domain A | 14200_8#1 | ND | 14623_8#85 | 23 | ND | - |
| | 1387582 | A > AT | frameshift | p.Ile160fs | domain A | 14623_4#78 | ND | 14323_1#84 | 66 | ND | - |
| | 1387954 | TC > T | frameshift | p.Glu36fs | domain A | 14623_3#1 | ND | 8524_5#62 | 45 | ND | - |

| | | | | | | | | | | | |
|---|---|---|---|---|---|---|---|---|---|---|---|
| *vraA* (SAOUHSC_ 00557) | 601292 | C > T | stop gained | p.Gln13* | domain A | 14208_8#78 | 1.5 | 14200_6#58 | 76 | 0.064 | *mprF* p.Thr345Ala |
| | 601292 | C > T | stop gained | p.Gln13* | domain A | 14623_8#74 | 2 | 14200_6#58 | 71 | 0.064 | *mprF* p.Thr345Ala |
| | 601292 | C > T | stop gained | p.Gln13* | domain A | 8447_7#54 | 1.5 | 14200_6#58 | 60 | 0.064 | *mprF* p.Thr345Ala |
| | 601616 | C > T | stop gained | p.Gln121* | domain A | 14200_8#40 | 0.25 | 8525_1#80 | 84 | 0.38 | - |
| | 601629 | AT > A | frameshift | p.Asn127fs | domain A | 14412_8#43 | ND | 8490_3#77 | 546 | ND | - |
| | 601629 | AT > A | frameshift | p.Asn127fs | domain A | 14448_4#53 | ND | 8490_3#77 | 575 | ND | - |
| | 601809 | T > A | stop gained | p.Leu185* | domain A | 14623_3#35 | 0.38 | 14200_8#1 | 69 | 0.38 | - |
| | 601809 | T > A | stop gained | p.Leu185* | domain A | 14623_3#89 | 0.25 | 14200_8#1 | 69 | 0.38 | - |
| | 601844 | C > T | stop gained | p.Gln197* | domain A | 14623_6#73 | ND | 14355_2#73 | 202 | ND | - |
| | 602186 | G > T | stop gained | p.Gly311* | domain A | 14412_8#68 | 0.5 | 8490_3#55 | 65 | 0.38 | - |
| | 602186 | G > T | stop gained | p.Gly311* | domain A | 14623_6#28 | 0.38 | 8490_3#55 | 65 | 0.38 | - |
| | 602186 | G > T | stop gained | p.Gly311* | domain A | 14623_7#77 | ND | 8490_3#55 | 68 | ND | - |
| | 602286 | TAA > T | frameshift | p.Lys345fs | domain A | 14355_1#1 | ND | 14355_2#9 | 532 | ND | - |
| | 602286 | TAA > T | frameshift | p.Lys345fs | domain A | 14355_1#3 | ND | 14355_2#9 | 500 | ND | - |
| | 602286 | TAA > T | frameshift | p.Lys345fs | domain A | 14623_4#87 | ND | 14355_2#9 | 500 | ND | - |
| | 602286 | TAA > T | frameshift | p.Lys345fs | domain A | 8447_7#44 | ND | 14355_2#9 | 500 | ND | - |
| | 602286 | TAA > T | frameshift | p.Lys345fs | domain A | 8447_7#45 | ND | 14355_2#9 | 500 | ND | - |
| | 602286 | TAA > T | frameshift | p.Lys345fs | domain A | 8490_3#61 | ND | 14355_2#9 | 500 | ND | - |
| | 602505 | T > A | stop gained | **p.Leu417*** | domain B | 14208_8#20 | ND | 8525_1#72 | 25 | ND | - |
| | 602505 | T > A | stop gained | **p.Leu417*** | domain B | 14623_7#13 | ND | none found | NA | ND | - |

[1]Mutations in bold were found to be homoplasic. [2]See information about vraA and pstS protein domains in the footnote of Supplementary Table 5. [3]Daptomycin MICs were determined using E-test. For each isolate, the genetically closest isolate was selected. Not all isolates

were selected for daptomycin susceptibility testing. Of all detected loss-of-function mutations, isolates were selected if the carried a nonsense (stop gained) mutation or homoplasic frameshift mutation, and had a genetically related isolate below 100 SNPs. If multiple isolates carrying the same mutation formed a monophyletic clade, then no more than two isolates with such mutation were selected for testing. *vraA* LOF mutations did not led to a measurable increase in daptomycin MIC with the exception of isolates carrying also a well-known daptomycin-resistance conferring mutation (*mprF* p.Thr345Ala). See Supplementary Table 7 for a comprehensive list of daptomycin-resistance conferring mutations.

Supplementary Table 7. Protein-altering mutations in the AgrCA two-component system identified in colonising isolates of the same host

| Gene (locus tag) | Chr. position | Nucleotide change | Annotation | Amino acid change | Protein domain | Patient id (isolate id) | Selected for testing (criterium)* |
|---|---|---|---|---|---|---|---|
| *agrC* (SAOUHSC_02264) | 2094797 | TTTA > T | DID | p.Ile52del | TMD 2 | chow2017-CD140408 (17059_1#56) | not available |
| | 2094952 | C > CT | frameshift | p.Val103fs | TMD 4 | tong2015-T197 (4950_3#3) | not available |
| | 2094974 | AT > A | frameshift | p.Ser109fs | TMD 4 | coll2017-675 (14412_8#81) | selected (TMD, frameshift) |
| | 2095085 | A > G | missense | p.Thr145Ala | TMD 5 | coll2017-308 (14448_2#30) | not selected |
| | 2095306 | G > A | missense | p.Met218Ile | DHp (HK) | coll2017-191 (14355_2#69) | not selected |
| | 2095340 | AC > A | frameshift | p.Thr230fs | DHp (HK) | coll2017-624 (14355_1#90) | not selected |
| | 2095399 | TA > T | frameshift | p.Asn251fs | DHp (HK) | coll2017-257 (8524_5#5) | not selected |
| | 2095409 | G > A | missense | p.Val253Ile | DHp (HK) | coll2017-416 (14324_8#66) | selected (DHp, missense) |
| | 2095435 | G > A | missense | p.Met261Ile | DHp (HK) | coll2017-79 (14623_2#2) | not selected |
| | 2095501 | GA > G | frameshift | p.Ile285fs | DHp (HK) | coll2017-273 (8524_5#30) | not selected |
| | 2095517 | C > T | stop gained | p.Gln289* | DHp (HK) | coll2017-571 (14623_3#73) | not selected |
| | 2095556 | G > T | stop gained | p.Glu302* | DHp (HK) | coll2017-764 (14324_8#39) | selected (DHp, stop codon) |
| | 2095556 | G > T | stop gained | p.Glu302* | DHp (HK) | coll2017-571 (14623_3#73) | not selected |
| | 2095592 | AGTC > A | DID | p.Arg315del | CA (HK) | coll2017-132 (14448_1#76) | selected (CA, inframe deletion) |
| | 2095605 | G > T | missense | p.Gly318Val | CA (HK) | coll2017-1232 (14623_4#15) | not selected |

| | | | | | | | |
|---|---|---|---|---|---|---|---|
| | 2095668 | C > T | missense | p.Ala339Val | CA (HK) | auguet2016-P_0130 (ERR1040796) | not available |
| | 2095711 | T > TA | frameshift | p.Cys355fs | CA (HK) | coll2017-1263 (14623_7#55) | selected (CA, frameshift) |
| | 2095724 | GA > G | frameshift | p.Asp358fs | CA (HK) | chow2017-CD141173 (16870_4#49) | not available |
| | 2095785 | G > A | missense | p.Gly378Asp | CA (HK) | coll2017-547 (14200_8#21) | selected (CA, missense) |
| | 2095808 | G > T | stop gained | p.Glu386* | CA (HK) | price2013-X077 (SRR2054058) | not available |
| | 2095836 | T > A | stop gained | p.Leu395* | CA (HK) | price2016-H211 (SRR3730915) | not available |
| *agrA* (SAOUHSC_02265) | 2096082 | A > G | missense | p.Tyr26Cys | RR | coll2017-450 (14200_6#83) | not selected |
| | 2096118 | G > A | missense | p.Gly38Asp | RR | coll2017-719 (14355_2#54) | selected (RR, missense) |
| | 2096130 | G > A | missense | p.Gly42Asp | RR | price2016-H213 (SRR3728573) | not available |
| | 2096292 | CT > C | frameshift | p.Ala97fs | ID | coll2017-364 (14412_8#75) | selected (ID, frameshift) |
| | 2096358 | G > A | missense | p.Gly118Asp | DNA BD | price2016-3449573 (SRR3729415) | not available |
| | 2096358 | G > A | missense | p.Gly118Asp | DNA BD | coll2017-1295 (14412_8#14) | selected (DNA BD, missense) |
| | 2096390 | A > G | missense | p.Ile129Val | DNA BD | coll2017-95 (8524_3#54) | not selected |
| | 2096395 | GT > G | frameshift | p.Phe132fs | DNA BD | coll2017-261 (14355_1#24) | not selected |
| | 2096395 | G > GT | frameshift | p.Glu133fs | DNA BD | coll2017-254 (14200_8#72) | selected (DNA BD, frameshift) |
| | 2096395 | G > GT | frameshift | p.Glu133fs | DNA BD | coll2017-503 (14623_8#17) | not selected |
| | 2096395 | G > GT | frameshift | p.Glu133fs | DNA BD | coll2017-377 (14623_5#51) | not selected |
| | 2096418 | C > A | missense | p.Ser138Tyr | DNA BD | price2016-H153 (SRR3728921) | not available |
| | 2096447 | C > T | missense | p.Arg148Cys | DNA BD | price2013-X168 (SRR2054034) | not available |

| | | | | | | | |
|---|---|---|---|---|---|---|---|
| | 2096462 | T > C | missense | p.Tyr153His | DNA BD | price2016-H122 (SRR3730764) | not available |
| | 2096499 | G > A | missense | p.Arg165His | DNA BD | price2013-X054 (SRR2054010) | not available |
| | 2096562 | A > T | missense | p.Lys186Ile | DNA BD | coll2017-190 (8447_7#11) | selected (DNA BD, missense) |
| | 2096567 | C > T | stop gained | p.Arg188* | DNA BD | young2017-P042 (SRR5251255) | not available |
| | 2096582 | A > T | stop gained | p.Lys193* | DNA BD | price2016-H236 (SRR3728784) | not available |
| | 2096590 | 41nt > A | frameshift | p.Glu196fs | DNA BD | harrison2016-GR13 (13414_7#6) | not available |
| *hld-argB* intergenic region | 2093655 | C > T (-178) | - | - | - | chow2017-CD142136 (17150_1#50) | not available |
| | 2093784 | C > T (-49) | - | - | AgrA BS | coll2017-257 (14623_8#22) | selected (intergenic) |
| | 2093801 | C > T (-32) | - | - | AgrA BS | price2016-2754372 (SRR3731418) | not available |
| | 2093812 | G > A (-21) | - | - | AgrA BS | price2016-2633534 (SRR3731564) | not available |
| | 2093812 | G > A (-21) | - | - | AgrA BS | harkins2018-SS_094 (ERR1904013) | not available |

*Only isolates from internal collections (i.e. coll2017) were available for testing. One isolate representative of each type of mutation (missense, frameshift and stop gained) at domain were selected for further testing. Abbreviations: AgrA BS, AgrA Binding Site (AgrA tandem repeats); CA, Catalytic and ATP-binding subdomain of the Histidine Kinase domain; Dimerization and Histidine phosphotransfer subdomain of the Histidine Kinase domain; DID, disruptive inframe deletion; DNA BD, DNA Binding Domain; ID, Inter Domain (region between domains); RR, Response Regulator domain; TMD, transmembrane domain.

Supplementary Table 8. Collections with multiple sequenced colonising isolates available from the same individual identified after June 2019

| Study Accession (Publication) | Isolates used (out of available) | Isolates kept after QC | Sources of Isolation | Years of Isolation | Place of Isolation | Number of hosts | Median isolates per individual |
|---|---|---|---|---|---|---|---|
| PRJEB4140, PRJEB2076 & PRJEB2489[47] | 1865/1935 | 457 | Nasal, throat and axilla swab | 2008 | Ubon Ratchathani, Thailand | 34 | 14.5 (2-19.5) |
| PRJEB9390[48] | 717/1478 | 391 | Axilla, groin and nasal swab | 2014 - 2016 | Singapore | 186 | 2 (2-2) |
| PRJEB28206[49] | 318/784 | 282 | Nasal swabs | 2018 - 2019 | Cambridge, UK | 141 | 2 (2-2) |
| PRJDB5246[50] | 124/242 | 60 | Cheek skin | 2010 - 2014 | Chiba, Japan | 30 | 2 (2-2) |
| PRJEB33854[51] | 35/37 | 23 | Nasal swab | 2016 | Westphalia, Germany | 6 | 3 (3-4.5) |
| PRJEB40888[52] | 103/165 | 60 | Axilla, groin and nasal swab | 2012-2014 | Australia | 18 | 2.5 (2-3.75) |
| PRJEB43023[53] | 154/155 | 76 | Nasal swabs | 2012-2014 | Denmark | 14 | 5 (2-7.75) |
| PRJNA590514[54] | 254/313 | 216 | Groin and nasal swab | 2017 - 2018 | Maryland, USA | 74 | 3 (2-3.75) |
| PRJNA587530[55] | 199/244 | 151 | Nasal, throat, groin, and perianal swabs | 2015 | Georgia, USA | 37 | 3 (2-5) |
| PRJNA595570[56] | 96/127 | 40 | Nasal, hands, throat, groin, axilla, and perianal swabs | 2015 - 2016 | Chicago, Illinois, USA | 18 | 2 (2-2) |
| PRJNA530184[57] | 191/267 | 115 | Nasal, throat and groin swabs | 2016 - 2017 | Chicago, Illinois, USA | 47 | 2 (2-3) |
| PRJNA638400[58] | 50/220 | 38 | Nasal, throat and groin swabs | 2007 - 2017 | Midwest, USA | 16 | 2 (2-3) |
| PRJNA715375, PRJNA715649 & PRJNA816913[59] | 382/1531 | 112 | Nasal swab | 2017-2018 | Mexico City, Mexico | 19 | 5 (3-6.5) |
| PRJNA685142[60] | 97/140 | 68 | Pooled four-site surveillance swab | 2017-2018 | New York City, USA | 26 | 2 (2-3) |
| PRJNA918392[61] | 3675/3818 | 2001 | Nasal, throat and groin swabs | Not specified | Pennsylvania, USA | 136 | 11 (7.75-20.25) |

| | | | | | | | |
|---|---|---|---|---|---|---|---|
| Total | 8260/11456 | 4090 | | | | 802 | 2 (2-4) |

See footnote of Supplementary Table 1 for legend information.

Supplementary Figure 1 Selection criteria used to identify collections of multiple colonising isolates sequenced per host

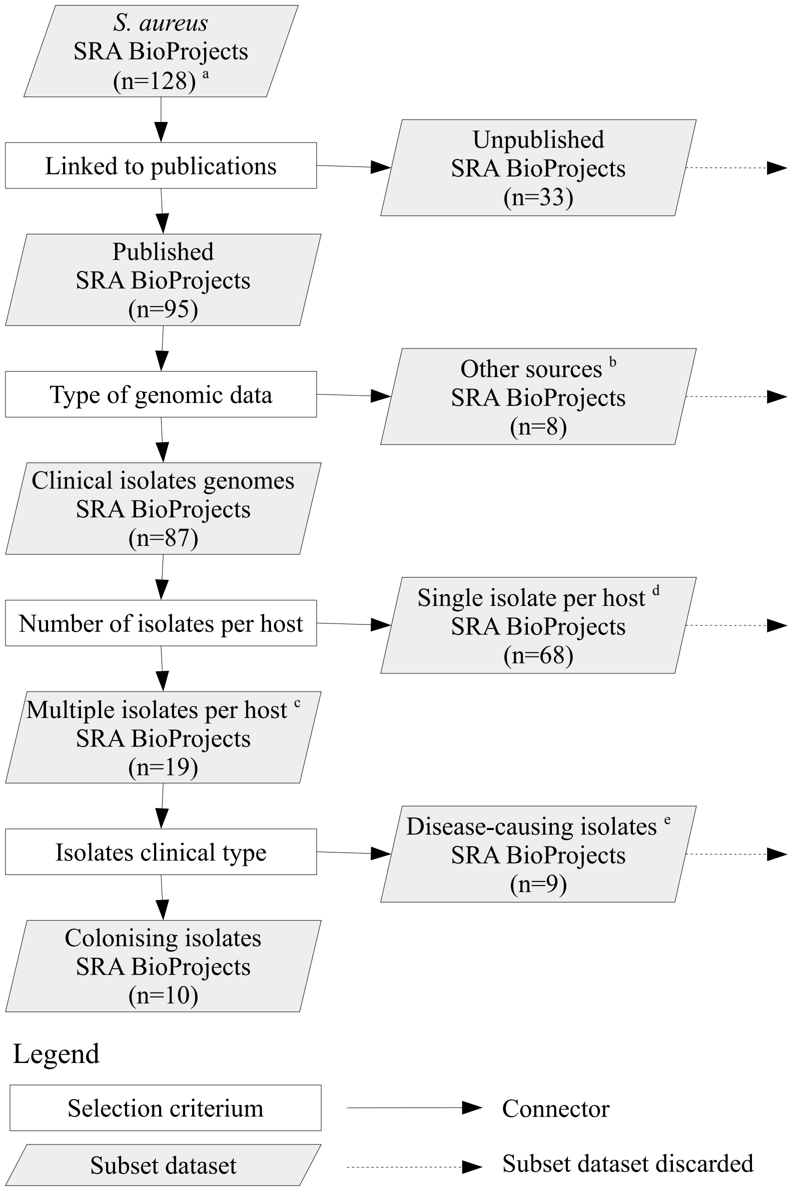

A total of 10 collections (SRA BioProjects) with multiple carriage isolates sequenced from the same host were identified after querying systematically for all *S. aureus* Short Read Archive (SRA) genomic data published to date (searched done by June 2019). [a] BioProjects with less than 70 *S. aureus* isolate genomes were discarded. [b] BioProjects including genomic data other than *S. aureus* clinical isolate genomes; that is, animal strains, mutagenesis experiments, RNAseq or microbiomes were discarded. [c] Host identifiers also had to be available in metadata to link isolates to their corresponding host. [d] BioProjects with multiple isolates per host but without host identifiers could not be included. [e] Datasets with multiple disease-causing isolates per host, or datasets with both colonising and disease-causing isolates per host but with only a single colonising isolate per host, were discarded.

Supplementary Figure 2 Genetic diversity between colonising isolates of the same host

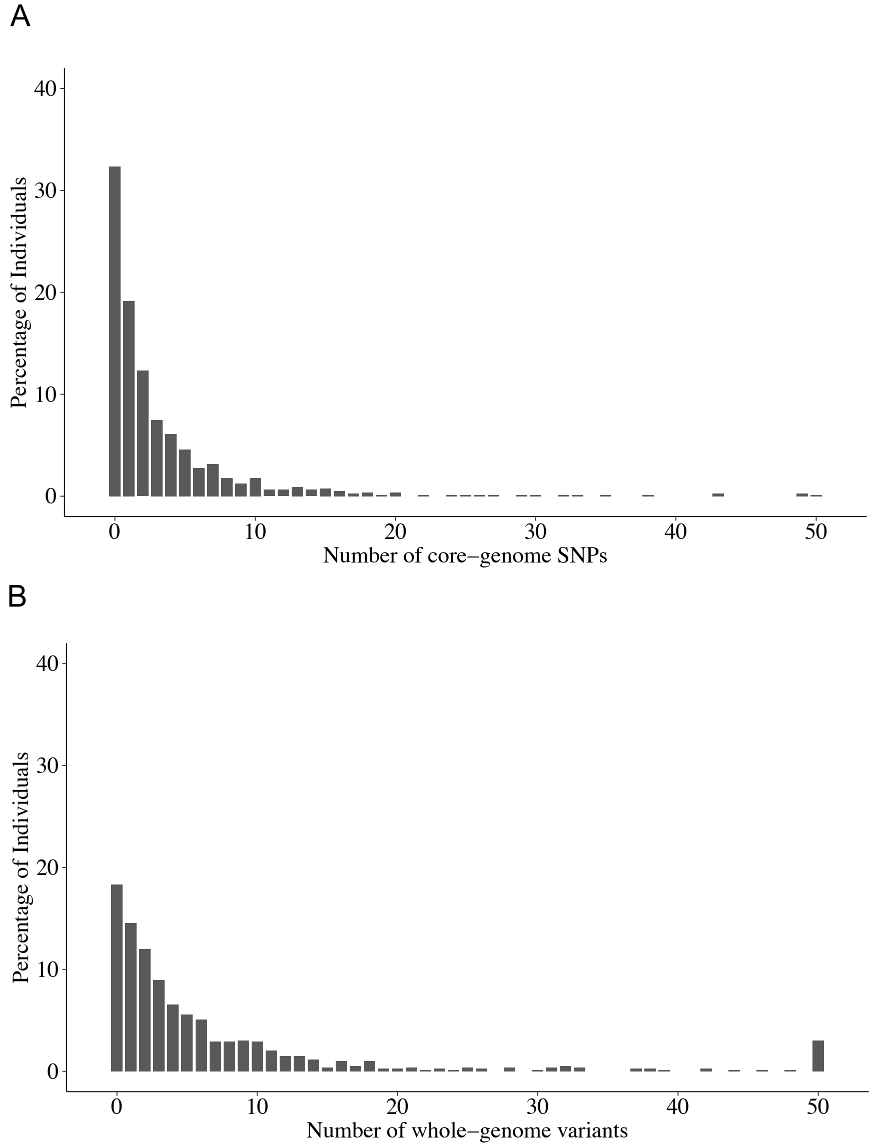

A. Genetic diversity measured as the maximum number of core-genome SNPs observed between isolates of the same individual. B. Genetic diversity measured as the total number of different genetic variants (SNPs and indels) in the whole genome observed across all isolates from the same individual. In the y-axis, percentage of individuals (out of total of 791) having the number of genetic variants specified in the x-axis.

Supplementary Figure 3. Density of mutations attributable to recombination

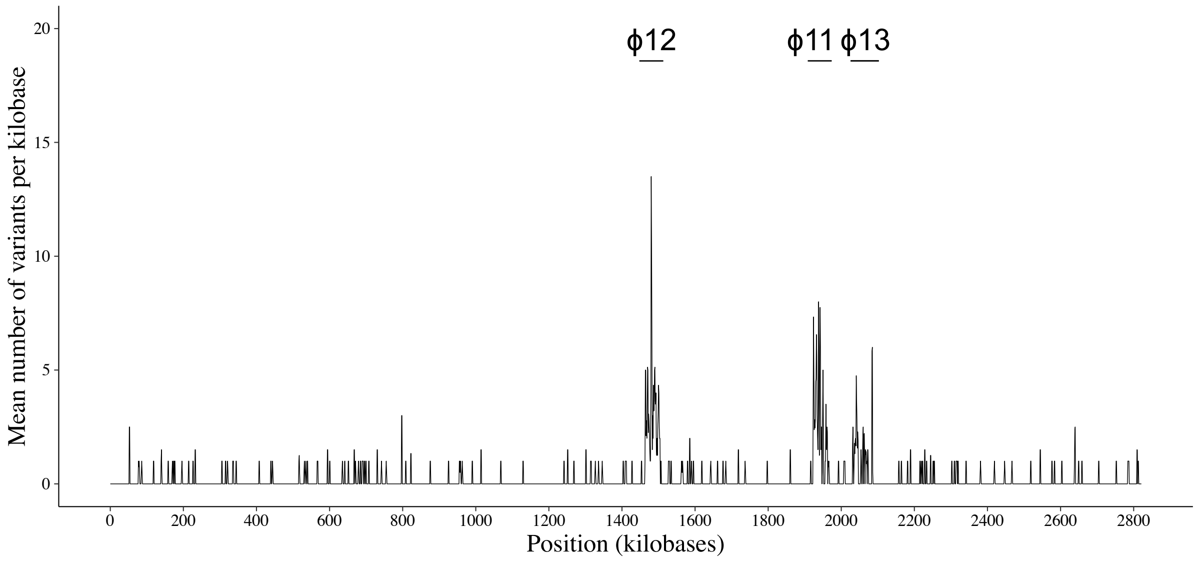

A sliding window of 2 kb was used to calculate the density of mutations attributable to recombination. The mean number of variants per kilobase is plotted along the NCTC8325 reference genome. Most recombination is concentrated in three regions of the genome annotated as prophages phi 11 (AF424781.1), phi 12 (AF424782.1) and phi 13 (AF424783.1).

Supplementary Figure 4 Growth curves of S. aureus nasD and ureG knock-out mutants under different nitrogen sources

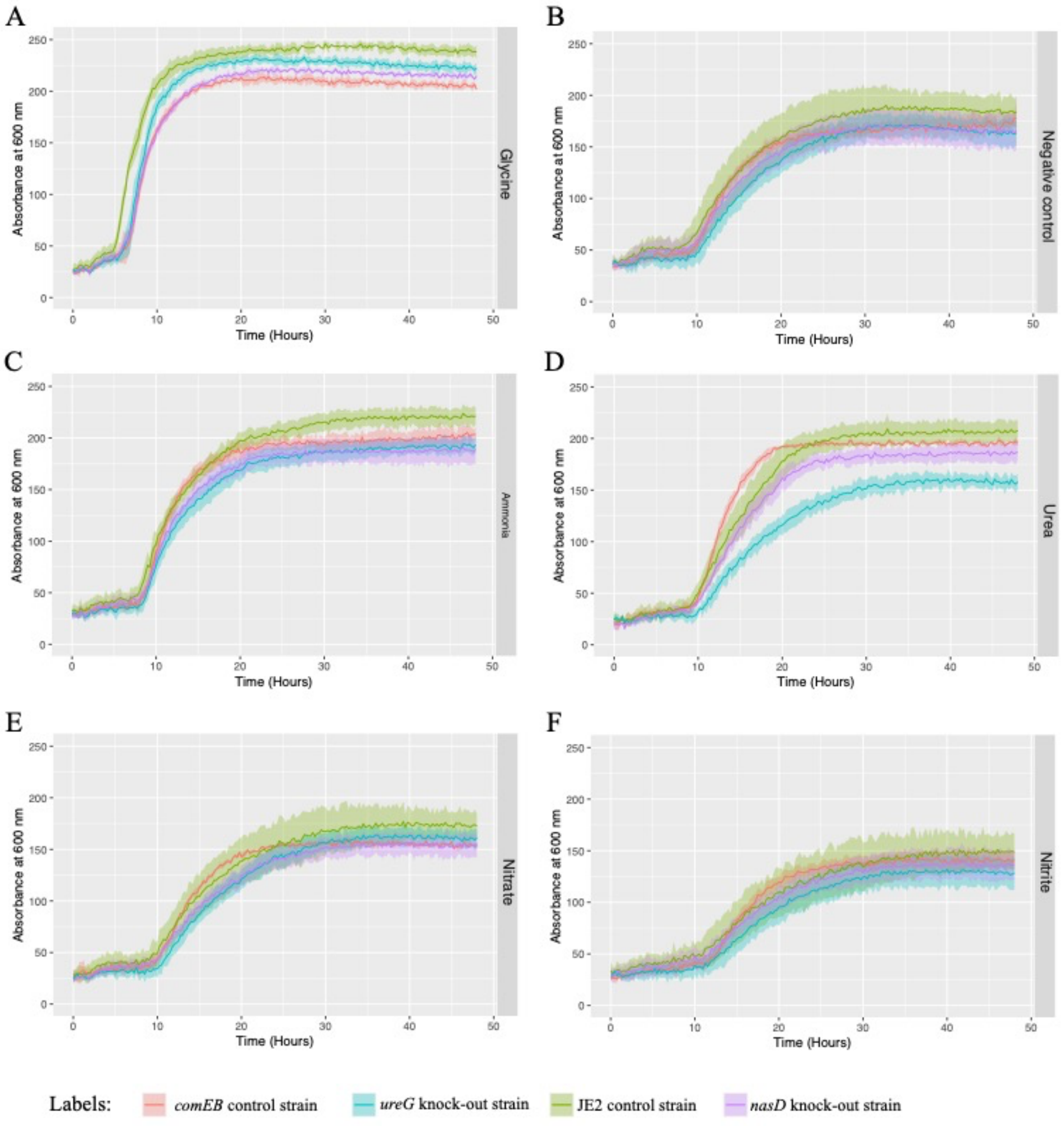

Growth curves of *S. aureus nasD/nirB*, *ureG* and *comEB* knock-out mutants; and JE control strain under the following nitrogen sources: glycine (panel A), negative control well (B), ammonia (C), urea (D), nitrate (E) and nitrite (F). Coloured lines represent mean OD600 calculated across three replicates, and shaded coloured regions the standard deviation.

Supplementary Figure 5. Growth curves of S. aureus *nasD*(*nirB*) mutants under different nitrogen sources

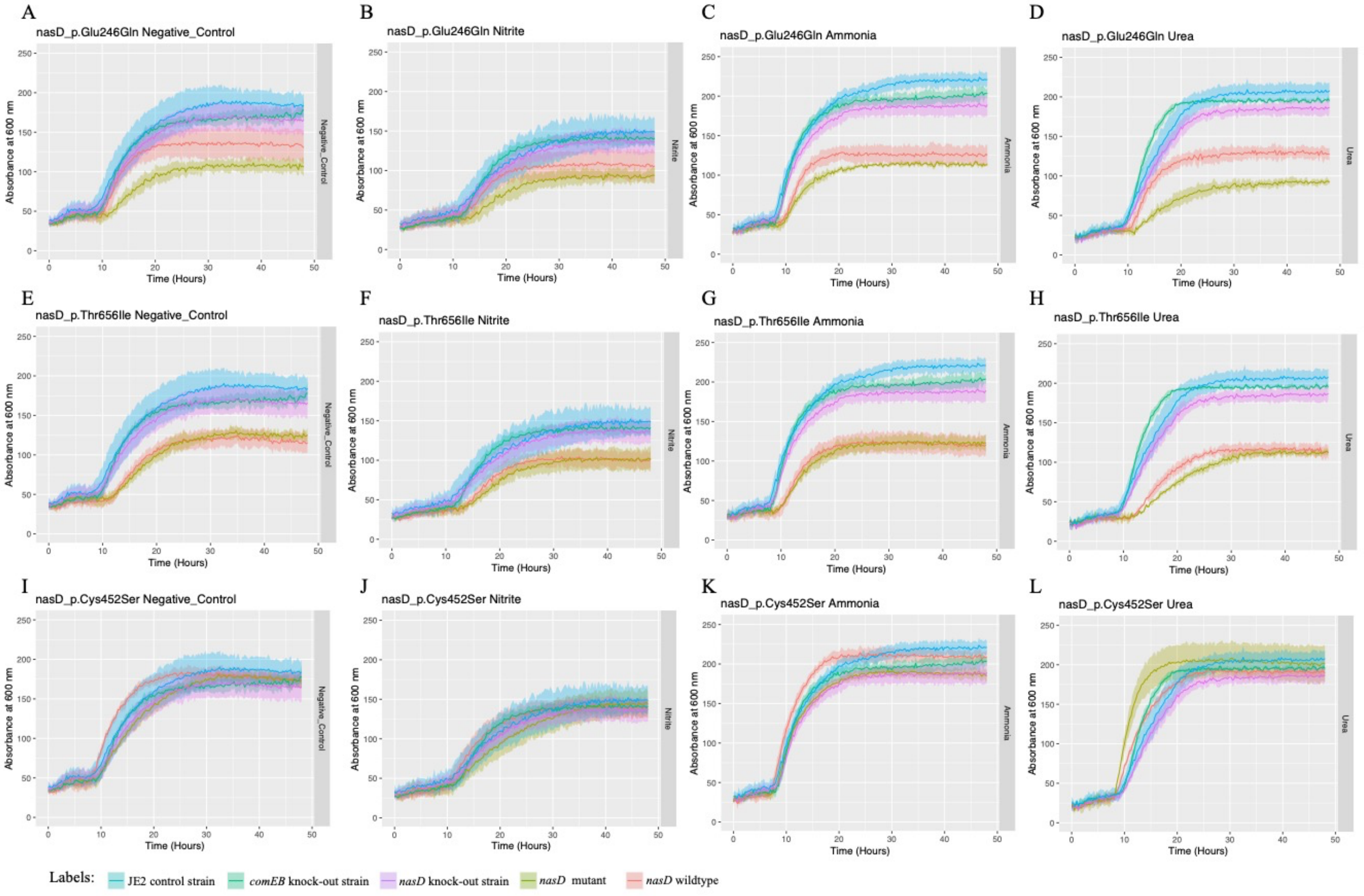

Growth curves of *S. aureus nasD/nirB* mutants, wildtypes (i.e. quasi-isogenic isolate lacking the *nasD/nirB* mutation from the same host), *nasD/nirB* knock-out, *comEB* knock-out (control) and JE (control) strain under the following nitrogen sources: negative control well, nitrite, ammonia, and urea. Coloured lines represent mean OD600 calculated across three replicates, and shaded coloured regions the standard deviation.

Supplementary Figure 6. Protein-altering mutations detected in known antibiotic targets.

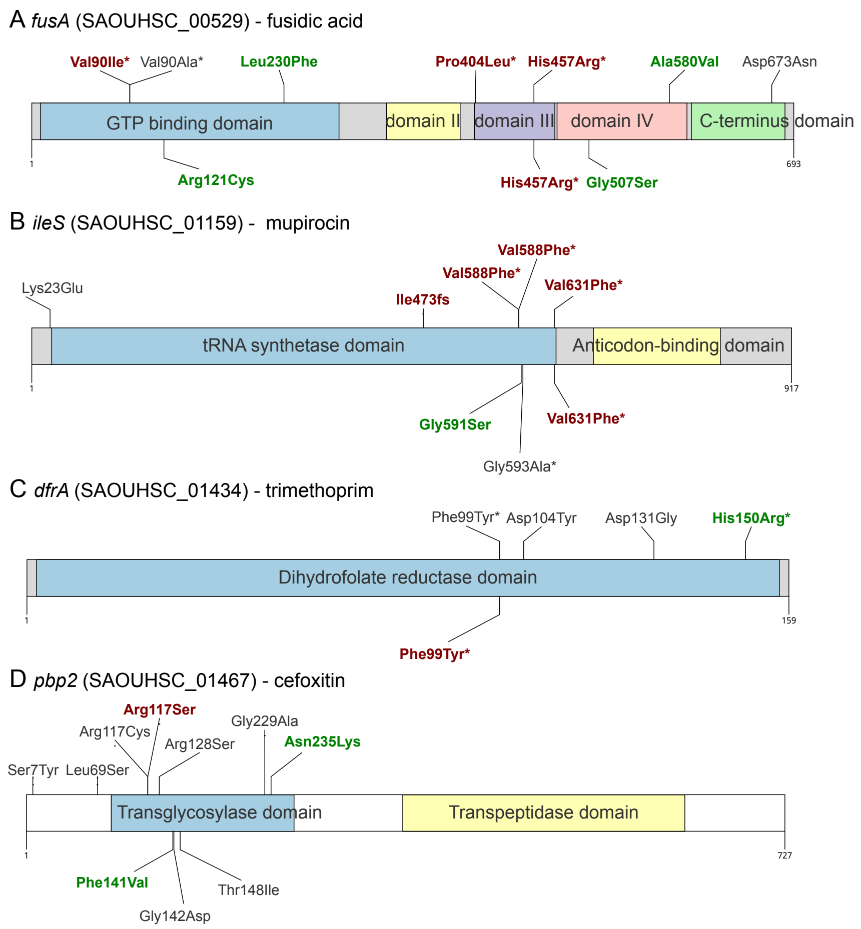

The proteins and protein domains of genes encoding antibiotics targets are shown. Mutations linked to a decreased susceptibility (increase in MIC) to their cognate antibiotic are coloured in red, while mutations in susceptible isolates are shown in green. Non-coloured mutations were found in external isolates that could not be tested. Asterisks indicate mutations reported to confer antibiotic resistance. (A) Elongation factor G, the target of fusidic acid, encoded by *fusA*. (B) Isoleucyl-tRNA synthetase, the target of mupirocin, encoded by *ileS*. (C) Dihydrofolate reductase, the target of trimethoprim, encoded by *dfrA*. (D) Penicillin-binding protein 2, target of beta-lactams, encoded by *pbp2*.

Supplementary Figure 7. Growth curves of *pstS* and *vraA* mutant and wildtype S. aureus clinical isolates under daptomycin exposure

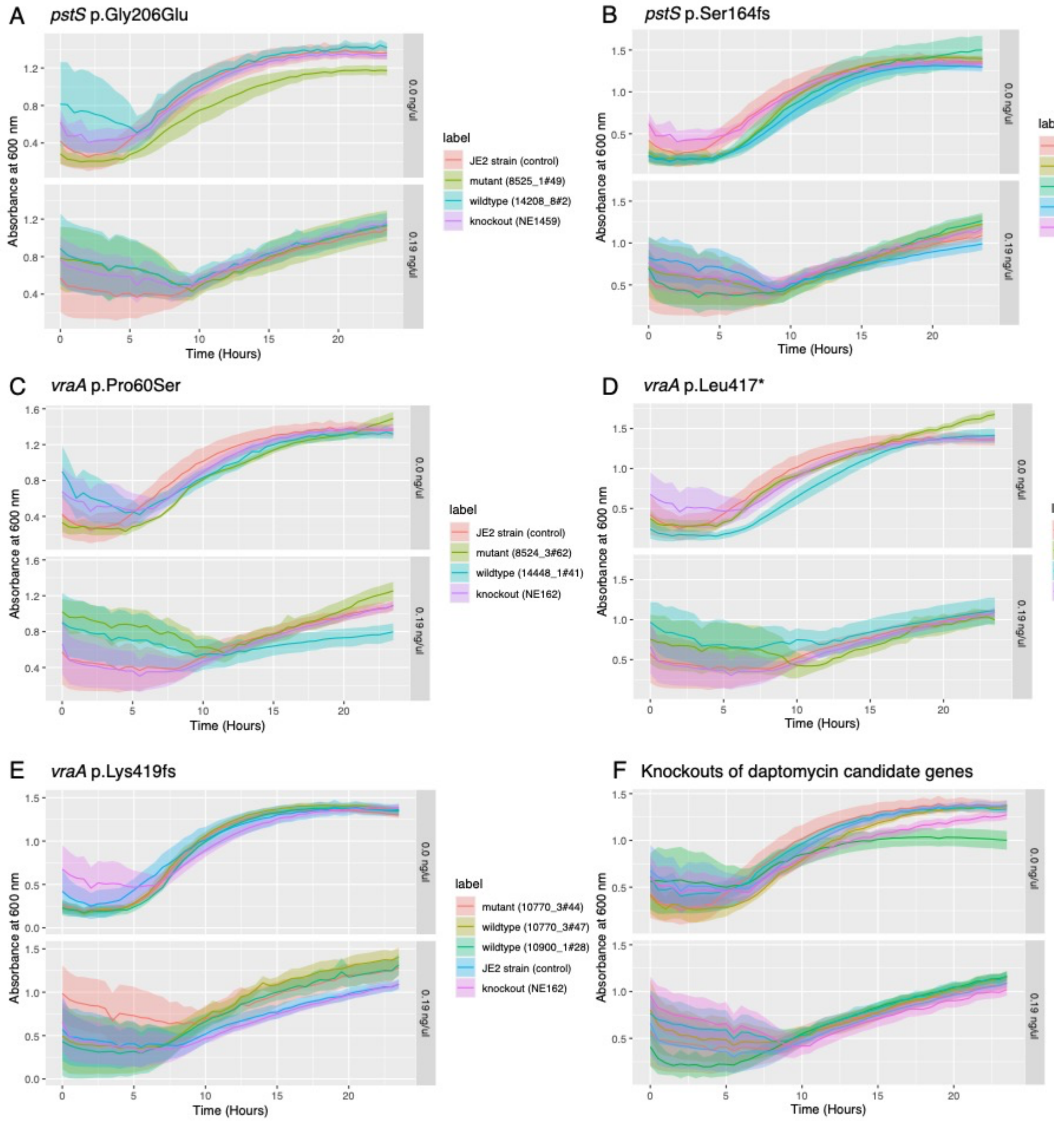

Panels A to E. Growth curves of *S. aureus* isolates carrying natural mutations in *pstS* and *vraA* genes in the absence and presence of sub-inhibitory concentration of daptomycin (0.19 µg/mL). As controls we grew wildtype *S. aureus* isolates from the same host lacking the investigated mutation, a transposon knockout of the investigated gene (*pstS* or *vraA*), and the *S. aureus* strain (JE2) used to build the transposon library. Panel F. Growth curves of *S. aureus* transposon knockouts in daptomycin-candidate genes. The values of absorbance at 600 nm are plotted in the y-axis as a function of time (in hours). Daptomycin concentration is indicated at the right edge of each plot.

Supplementary Figure 8. Loci enriched for protein-altering mutations in colonising isolates of the extended dataset

A CDS

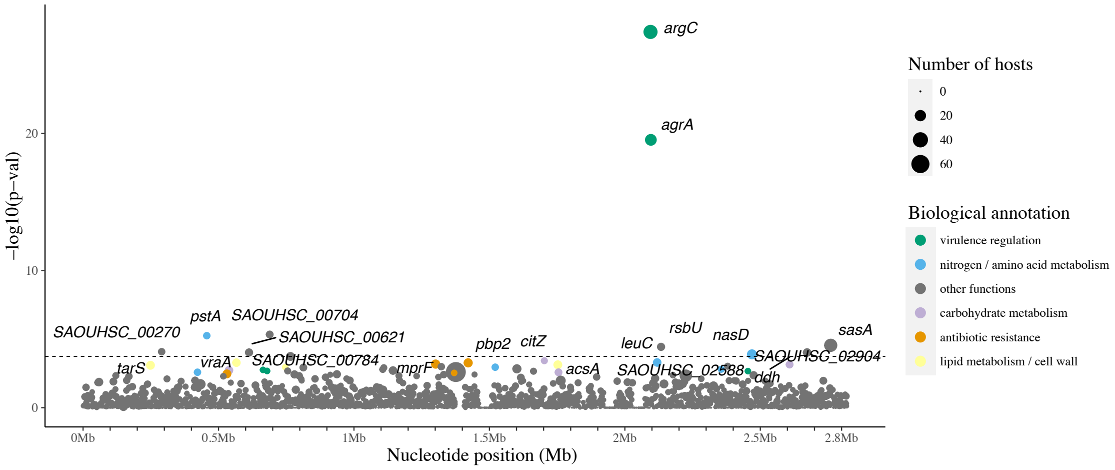

B Transcription units (operons)

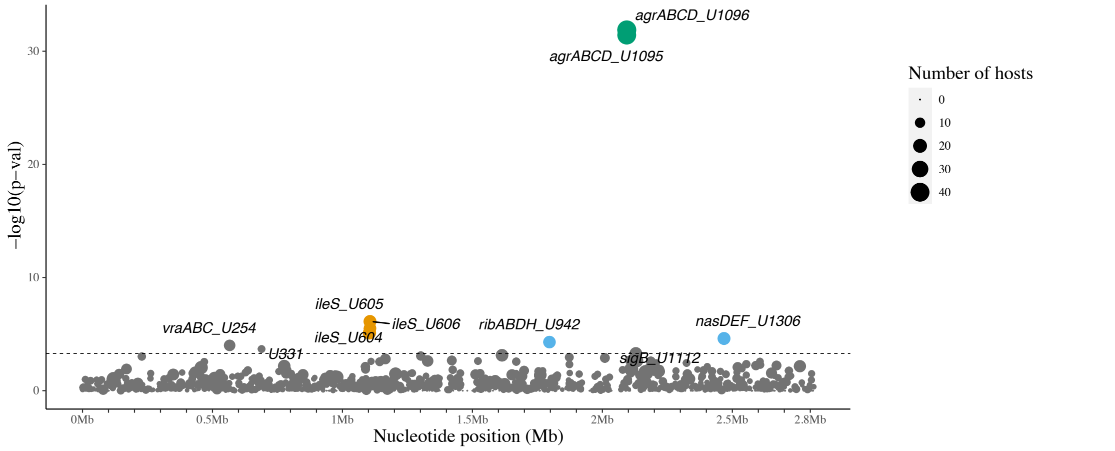

C Metabolic sub-modules

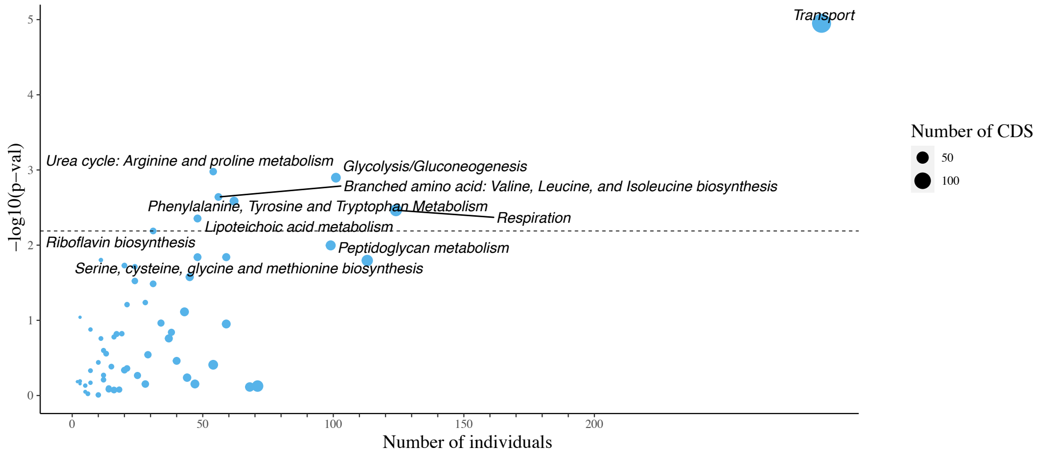

See footnote of Figure 2 for legend information.
